# Bioenergetic costs and the evolution of noise regulation by microRNAs

**DOI:** 10.1101/2023.03.28.534633

**Authors:** Efe Ilker, Michael Hinczewski

**Affiliations:** Max Planck Institute for the Physics of Complex Systems, 01187 Dresden, Germany; Department of Physics, Case Western Reserve University, Cleveland OH 44106

## Abstract

Noise control, together with other regulatory functions facilitated by microRNAs (miRNAs), is believed to have played important roles in the evolution of multicellular eukaryotic organisms. miRNAs can dampen protein fluctuations via enhanced degradation of mRNAs, but this requires compensation by increased mRNA transcription to maintain the same expression levels. The overall mechanism is metabolically expensive, leading to questions about how it might have evolved in the first place. We develop a stochastic model of miRNA noise regulation, coupled with a detailed analysis of the associated metabolic costs. Additionally we calculate binding free energies for a range of miRNA seeds, the short sequences which govern target recognition. We argue that natural selection may have fine-tuned the Michaelis-Menten constant *K*_M_ describing miRNA-mRNA affinity, and show supporting evidence from analysis of experimental data. *K*_M_ is constrained by seed length, and optimal noise control (minimum protein variance at a given energy cost) is achievable for seeds of 6-7 nucleotides in length, the most commonly observed types. Moreover, at optimality the degree of noise reduction approaches the theoretical bound set by the Wiener-Kolmogorov linear filter. The results illustrate how selective pressure toward energy efficiency has potentially shaped a crucial regulatory pathway in eukaryotes.

---

Nonequilibrium processes within living systems exact a high price: the constant maintenance of fuel molecules and raw materials at sufficient concentrations to provide thermodynamic driving potentials for biological function [1]. Optimizing that function with respect to thermodynamic costs is a factor constraining evolution, and would have been particularly important at the very earliest stages of life where the metabolic chemistry responsible for maintaining those potentials was necessarily primitive and relatively inefficient. Yet thermodynamic costs are not the only factor that matters, and biology is full of counter-intuitively complex chemical mechanisms whose evolutionary predecessors, perhaps arising out of the randomness of genetic drift, may have consumed energy resources without any clear fitness benefit.

The discovery of microRNAs (miRNAs) along with their counterparts, small non-coding RNAs, raised many open questions about their functional purposes and evolution [2, 3]. These short endogenous RNAs, around 22 nucleotides (nt) in length, exist in many eukaryotic cells. They constitute the core of the RNA-induced silencing complex (RISC) that interacts with target messenger RNA (mRNA), leading to translational repression and the accelerated degradation of their target by a mechanism known as the RNA interference. One possible functional role for this interference, which is the focus of our work, is fine-tuning noise in protein populations by reducing the variance of protein copy numbers [4, 5], conferring robustness to cellular functions [6]. Such noise control, together with other regulatory functions facilitated by microRNAs, is believed to have played important roles in the evolution of complex multi-cellular life [7–10]. Yet it is a considerable expenditure of resources, similar to setting up a factory production line for a valuable good, funding gangs of thieves to constantly raid the factory, and compensating for losses by increasing the production rate. *So how would such a regulation scheme arise, and has evolution actually optimized it?* Using a combination of statistical physics, information theory, biochemistry, and population genetics, we arrive at some tentative answers to these questions.

Our results give insights into one peculiar feature of this system: why the job requires such short RNA molecules—the significance of the “micro” in microRNA. Specific interactions between the miRNA and its target in fact largely depend on only a 6-8 nt sequence known as the miRNA seed region, which forms Watson-Crick base pairs with a complementary sequence on the target mRNA. We argue that the observed length of the seed region lies in a metabolic sweet spot, giving just enough affinity between the miRNA and mRNA (measured by their Michaelis-Menten constant *K*_M_) to optimally control noise at a given level of energetic expenditure. Longer seed sequences (higher affinity, smaller *K*_M_) would increase rather than decrease noise. On the other hand, shorter seed sequences (lower affinity, larger *K*_M_), while allowing interactions with a wider range of targets, would also require significantly more energy to achieve the same level of control. By estimating these energy expenditures, we also show that effective noise control is costly enough to be under selection pressure in eukaryotic cells. A novel miRNA may initially appear during the course of evolution as a random product of genetic drift with non-optimal parameters, and then get gradually repurposed as a noise control mechanism that confers a fitness advantage. Once this starts to occur, our theory predicts that natural selection would hone *K*_M_ toward an optimal value.

## miRNA regulation as a noise filter for protein expression

We start our theoretical description by introducing the architecture of the miRNA-regulated system 𝒮_*R*_ (Fig. 1B) in comparison to the unregulated one 𝒮_0_ (Fig. 1A). Our description builds off the experimentally-validated model of Ref. [5], and takes the form of stochastic biochemical reaction networks with dynamics obeying linearized chemical Langevin (CL) equations [11, 12], as detailed in Supplementary Information (SI) Sec. S.I A. The CL results agree closely with available experimental data [5], and give us analytical expressions for correlation functions describing copy number fluctuations of chemical species in each system.

**FIG. 1.**
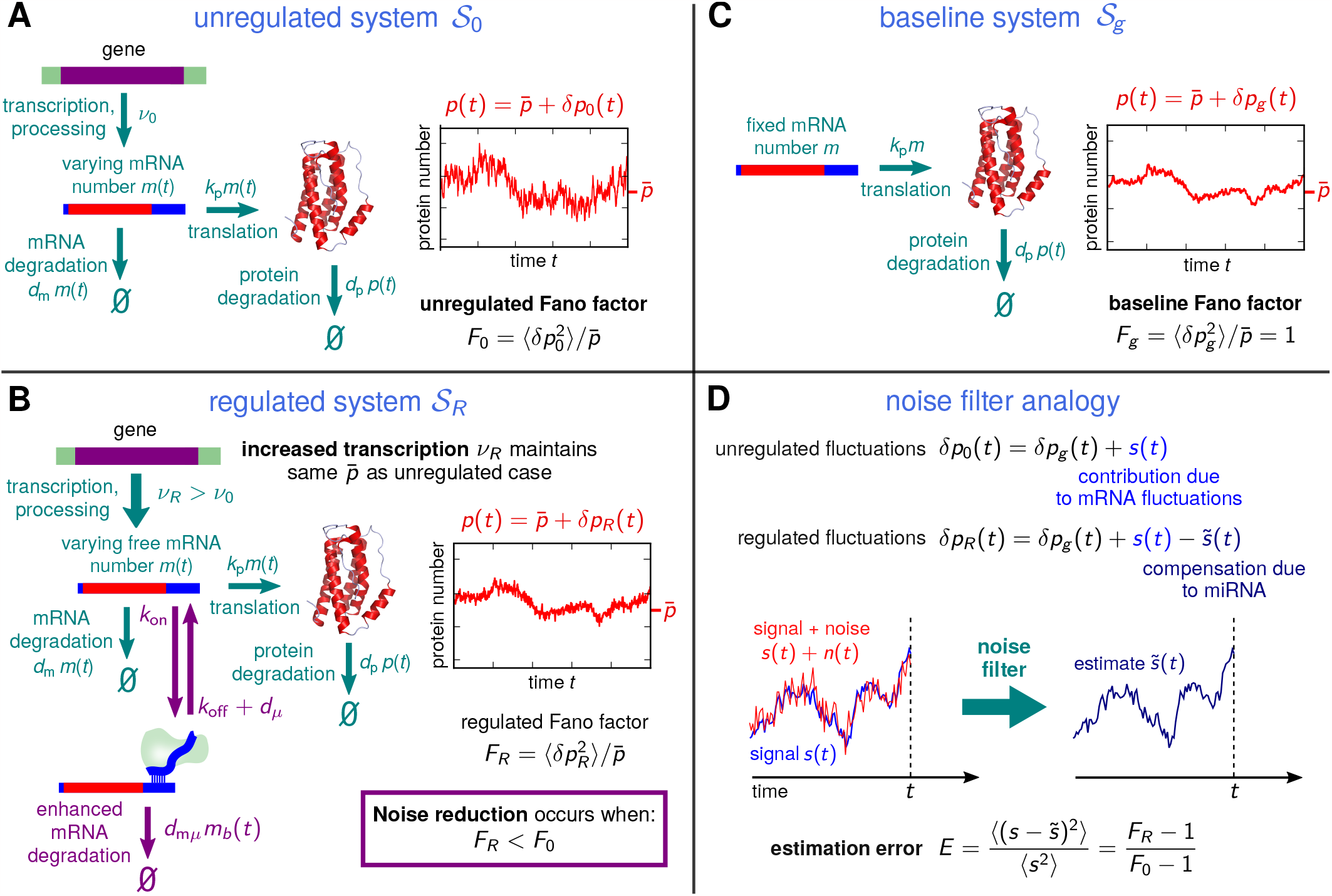
Overview of the miRNA noise regulation model: (**A**) In the unregulated system 𝒮_0_ with no miRNA, noise in protein numbers has contributions from both varying mRNA population and intrinsic noise in the translation process, leading to a Fano factor *F*_0_ (protein variance divided by the mean protein number 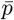). (**B**) For the regulated system 𝒮_*R*_, enhanced mRNA degradation due to targeting by miRNA can reduce protein noise, leading to a Fano factor *F*_*R*_ *< F*_0_. To compensate for the loss of mRNA and achieve the same 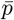, the transcription rate *ν*_*R*_ must be increased relative to its value *ν*_0_ in 𝒮_0_. (**C**) To set up the noise filter analogy, we introduce an imaginary basline system 𝒮_*g*_, which has fixed mRNA population and hence the protein noise is solely due to translation. The resulting Fano factor *F*_*g*_ = 1 is a lower bound on the two systems above: 1 ≤ *F*_0_, *F*_*R*_. (**D**) Decomposing the unregulated/regulated protein fluctuations *δp*_0_(*t*) and *δp*_*R*_(*t*) into a baseline contribution *δp*_*g*_ (*t*) and an additional contribution allows us to define the signal *s*(*t*) and estimate 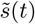 in the noise filter analogy. Their normalized mean-squared difference *E* is the estimation error of the filter, which can be expressed in terms of *F*_*R*_ and *F*_0_.

The fluctuating output numbers are denoted as 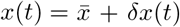 where *δx*(*t*) is the variation from the mean level 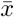. The main species of interest are total (bound + unbound) miRNA, mRNA and protein copy numbers, denoted by *x* = *μ*_tot_, *m* or *p* respectively. In the unregulated system 𝒮_0_ (Fig. 1A) we have transcription from a gene at rate *ν*_0_ producing an mRNA population *m*(*t*), which is then translated with rate constant *k*_*p*_ to a produce a protein population *p*(*t*). The mRNA and proteins are degraded with rate constants *d*_*m*_ and *d*_*p*_ respectively.

To meaningfully compare 𝒮_*R*_ to 𝒮_0_, we assume parameter sets such that the mean protein output 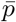 is the same for both regulated and unregulated systems. This maintains the functional effectiveness of the protein in the regulated system, but with the potential added benefit of noise reduction. Indeed miRNA are often upregulated along with their target mRNA via feedforward loops [13, 14]. For 𝒮_*R*_ (Fig. 1B) there is an additional RNA interference mechanism: free miRNAs with population *μ*(*t*) can bind to the mRNA with rate constant *k*_on_ to form a bound complex with population *m*_*b*_(*t*). Considering a single miRNA binding site on the target mRNA, the total number of miRNA is *μ*_tot_(*t*) = *μ*(*t*)+*m*_*b*_(*t*). The mRNA in the complex has an enhanced degradation rate constants *d*_*mμ*_ *> d*_*m*_ relative to the regular mRNA value *d*_*m*_. The miRNA unbinds with rate constant *k*_off_, and degrades with *d*_*μ*_. For simplicity we assume the miRNA degradation rate constant *d*_*μ*_ is the same when both bound and unbound to mRNA [5]. miRNA-mRNA affinity can be characterized through the Michaelis-Menten constant,

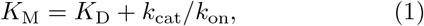

where the dissociation constant *K*_D_ = *k*_off_ */k*_on_ and *k*_cat_ = *d*_*mμ*_ + *d*_*μ*_ is an effective catalytic rate constant for the miRNA-catalyzed degradation reaction. *K*_M_ approximately relates *m*_*b*_(*t*) to *μ*_tot_(*t*) through *m*_*b*_(*t*) ≈ *m*(*t*)*μ*_tot_(*t*)*/*(*K*_M_ + *m*(*t*)). Note that experimental values of *K*_M_ and *K*_D_ are reported in units of concentration (molars) while our CL formalism uses copy numbers of chemical species. We assume a typical eukaryotic cell volume *V* = 2000 *μ*m^3^ to convert between concentrations and copy numbers as needed.

Because of interference from the miRNA, the transcription rate *ν*_*R*_ of mRNA in 𝒮_*R*_ must be larger than the rate *ν*_0_ in 𝒮_0_, if both systems maintain the same mean protein level 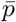. In the limit of no miRNA (*μ*_tot_ → 0) or vanishing affinity (*K*_M_ → ∞) the regulated system approaches the same behavior as the unregulated one, with *ν*_*R*_ → *ν*_0_. The strength of regulation can be characterized by a parameter *R* ≡ 1 − *ν*_0_*/ν*_*R*_, where *R* ranges between 0 (no regulation) to 1 (maximum possible regulation). For known miRNA-mediated regulation networks, *R* typically lies between 0.05-0.95 [15, 16].

To quantify the effect of miRNA on noise, we look at the Fano factor 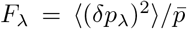 with *λ* = *R*, 0 labeling the system 𝒮_*λ*_ in which the quantity is calculated. Successful noise reduction implies that *F*_*R*_ *< F*_0_: for the same mean protein output level, there is less protein variance in the presence of miRNA. Qualitatively this arises because the miRNA system reduces the number of translated proteins per mRNA, on average by a factor of 1 − *R*, and hence decreases the susceptibility of translation to fluctuations in mRNA levels. When miRNA regulation is compensated for by transcriptional increase, it is thus possible to mitigate the propagation of noise from mRNA to protein numbers. However, there is a trade-off, because the stochasticity of miRNA-mRNA interactions, as well as fluctuating miRNA populations, also introduces noise into mRNA levels. This added noise can cancel out the protein noise reduction benefit in certain parameter regimes, for example when miRNA-mRNA affinities are high.

To understand the role of miRNA more precisely, it is helpful to use a noise filter analogy. In order to motivate this mathematical analogy, we define an imaginary baseline system 𝒮_*g*_ (Fig. 1C), where we have fixed the mRNA population at a constant level 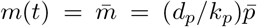, which agrees with the mean 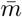 in 𝒮_*R*_ and 𝒮_0_ and hence gives the same 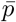. This removes the contribution of mRNA fluctuations to the noise, so 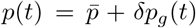, where *δp*_*g*_(*t*) are the ground-level (baseline) fluctuations that come from protein translation and degradation (and cannot be mitigated by RNA interference). As summarized in Fig. 1D, the protein fluctuations in both 𝒮_0_ and 𝒮_*R*_ can then be compared to this baseline. For 𝒮_0_ we write *δp*_0_(*t*) = *δp*_*g*_(*t*) + *s*(*t*), where *s*(*t*) represents the added noise due to varying mRNA population *m*(*t*). For 𝒮_*R*_ this added noise is partially compensated, 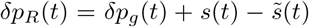. Both *s*(*t*) and 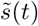 can be explicitly calculated using the CL formalism (SI Sec. S.II) and it turns out they are correlated: 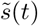 takes the form of a convolution,

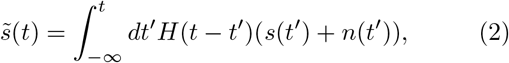

where *H*(*t*) and *n*(*t*) are functions of the biochemical parameters. The above equation has a clear noise filter interpretation (depicted schematically in Fig. 1D): *s*(*t*) is the “signal”, *n*(*t*) a “noise” that corrupts the signal, *H*(*t*) is a linear filter function that acts via convolution on the past history of the corrupted signal *s*(*t*) + *n*(*t*), and 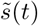 is the “estimate” of the signal. It turns out the problem of reducing protein noise via mRNA interference (making *F*_*R*_ as small as possible) is equivalent to making 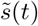 as close as possible to *s*(*t*). We can see this directly in the error of estimation, which is defined as 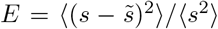. For our case, *E* can be expressed in terms of the Fano factors, *E* = (*F*_*R*_ − 1)*/*(*F*_0_ − 1). Fine-tuning miRNA parameters only affects *F*_*R*_, leaving *F*_0_ fixed, so *E* can be minimized by decreasing *F*_*R*_. Both *F*_*R*_ and *F*_0_ are bounded from below by the Fano factor *F*_*g*_ = 1 of the baseline system, so perfect filtering, *E* → 0, would correspond to *F*_*R*_ → 1. The noise filter interpretation, which has been earlier applied to a variety of other biological networks (for a review see Ref. [17]), has an important payoff which we will return to later: it allows us to find the conditions for optimal noise reduction, and calculate tighter bounds on *F*_*R*_, which in general will be greater than 1 at optimality.

## Bioenergetic costs of miRNA regulation

The second major component of our model is an estimate of the costs for miRNA regulation, which we adapt from experimental data on eukaryotic transcription energetics collected in Ref. [18]. In general this includes energy expenditures channeled to the synthesis of new molecules, as well as maintenance, the recycling/repair of molecules to maintain steady-state levels (i.e. assembling mRNA from existing nucleotides to counterbalance degradation). Eukaryotic cells typically have long enough generation (cell division) times *t*_*r*_, that maintenance is the dominant contribution to metabolic expenditures over a generation. Focusing on the maintenance costs, we can estimate the total metabolic consumption *C*_*T*_ (in units of phosphate [P] bonds hydrolyzed, namely ATP or ATP-equivalents) for the unregulated system per generation: *C*_*T*_ ≃ *t*_*r*_*M*_*ν*_, where *M*_*ν*_ = *ν*_0_*ϵ*_*m*_ is the consumption rate. Here *ν*_0_ is the mRNA transcription rate in 𝒮_0_, and *ϵ*_*m*_ is the energy cost in terms of P for assembling the mRNA (which will depend on the length of the transcript). For the regulated case, the total cost will be *C*_*T*_ + *δC*_*T*_, where the extra contribution *δC*_*T*_ = *t*_*r*_Δ*M*_*ν*_. The difference in consumption rate Δ*M*_*ν*_ = (*ν*_*R*_ − *ν*_0_)*ϵ*_*m*_ +*ν*_*μ*_*ϵ*_*μ*_. The first term accounts for the added costs of increased transcription to maintain the same 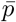, while the second term is the rate of miRNA assembly *ν*_*μ*_ times the cost *ϵ*_*μ*_ of that assembly in units of P (including potentially any related costs of the RISC complex). As shown in SI Sec. S.I C, Δ*M*_*ν*_ can be expressed in terms of the biochemical parameters of the system as

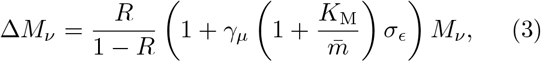

with *γ*_*μ*_ ≡ *d*_*μ*_*/d*_*mμ*_ and *σ*_*ϵ*_ ≡ *ϵ*_*μ*_*/ϵ*_*m*_. Based on experimental estimates (summarized in SI Table S1) we know that *γ*_*μ*_ ≪ 1 and *σ*_*ϵ*_ ≪ 1. Given the experimental range of *R* = 0.05−0.95, the term *R/*(1 − *R*) can vary by a factor of 361 between the smallest and largest observed regulation magnitudes, highlighting the strong dependence of Δ*M*_*ν*_ on *R*. The remaining terms in Eq. [3] encapsulate the modification to the costs due the parameters governing miRNA-mRNA interactions, particularly *K*_M_ and the degradation enhancement *d*_*mμ*_.

As discussed later in the section on evolutionary pressure, it is convenient to define a non-dimensional measure for the extra cost due to regulation: the extra cost as a fraction of the total metabolic expenditure of a cell per generation, *δC*_*T*_ */C*_*T*_. Here *C*_*T*_ ≃ *t*_*r*_*M*_tot_, where *M*_tot_ is the total maintenance ATP consumption rate. Based on the data from Ref. [18], *M*_tot_ approximately scales with cell volume, and we use a value *M*_tot_ = 3 × 10^11^ P/hr characteristic of eukaryotic cells. Thus we will report costs in terms of *δC*_*T*_ */C*_*T*_ = Δ*M*_*ν*_*/M*_tot_, with Δ*M*_*ν*_ given by Eq. [3].

Our model so far has assumed one mRNA target, but in general a single miRNA can target up to hundreds of mRNAs, which will also change the energetic costs of regulation. As a rough estimate of this multi-target scenario, we can assume similar biochemical parameters among different targets. This allows us to use the single-target theory but scaling up 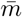 (and hence 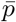) to reflect the total mRNA numbers when accounting for all targets. Note that the dependence of Δ*M*_*ν*_ on 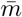 is non-trivial, since both *R* and *M*_*ν*_ depend on 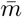. However when we demand a certain level of overall noise control (a specific value of *E*) the extra cost Δ*M*_*ν*_ will increase with 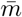 as the number of targets gets larger: the larger the system, the more expensive it is to control.

## Seeds of length 6 − 7 nt are most energetically efficient for noise reduction

With all the components of the model defined, we can now investigate the interplay between noise reduction and energetic costs. The error *E* (or equivalently the Fano factor *F*_*R*_) can be expressed as a function of Δ*M*_*ν*_, *γ*_*μ*_, *σ*_*ϵ*_, 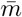 and *K*_M_ (details in the SI Secs. S.I B,C). For a given cost Δ*M*_*ν*_ and fixed degradation/energy parameters *γ*_*μ*_ and *σ*_*ϵ*_, we can ask what value of *K*_M_ minimizes *E. K*_M_ is an interesting tuning parameter because it is related to the binding strength between the miRNA and the mRNA, which in turn depends on the number of complementary interactions between the seed and the target region of the mRNA. From Eq. [1], *K*_M_ ≥ *K*_D_, and the dissociation constant *K*_D_ is related to the free energy of binding via Δ*G*^0^ = *k*_*B*_*T* ln(*K*_D_*/*[1M]). Longer seeds should allow for more negative Δ*G*^0^ (stronger binding) and hence smaller values of *K*_D_. This in turn gives access to smaller values of *K*_M_. For RNA interference systems, experimentally measured *K*_M_ values have ranged from comparable to *K*_D_ to about two orders of magnitude larger than *K*_D_ [19].

In Fig. 2A, we show the contour diagram of log_10_ *E* as a function of *K*_M_ and fractional metabolic cost *δC*_*T*_ */C*_*T*_, assuming a single mRNA target and using the experimentally-derived parameters of SI Table S1. The blue curve denotes the optimal value 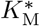, which achieves the minimum *E* for a given cost *δC*_*T*_ */C*_*T*_. The red contour line marks the value 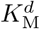 where *E* = 1, which corresponds to *F*_*R*_ = *F*_0_. This is the boundary between the noise control region to the right, where *E <* 1 (*F*_*R*_ *< F*_0_) and a “dud” region to the left, where *E >* 1 (*F*_*R*_ *> F*_0_). In the latter region regulation adds protein noise to the system rather than mitigating it which can provide an alternative role for some microRNA systems in triggering cell state transitions [20, 21]. For a fixed *δC*_*T*_ */C*_*T*_, as we scan *K*_M_ from small to large values, we cross from dud to noise control at 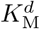, improve the filter performance until we reach 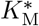, and then get progressively worse filtering for 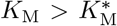. The different behaviors of the system with varying *K*_M_ reflect the tradeoff due to miRNA-mRNA affinity mentioned earlier in the noise filter discussion. The optimal affinity 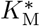 is a metabolic sweet spot between a regime where miRNA-mRNA interaction is too strong (small *K*_M_), leading to excessive added noise and the “dud” scenario, and a weak interaction regime (large *K*_M_) where the miRNA system cannot effectively dampen noise.

**FIG. 2.**
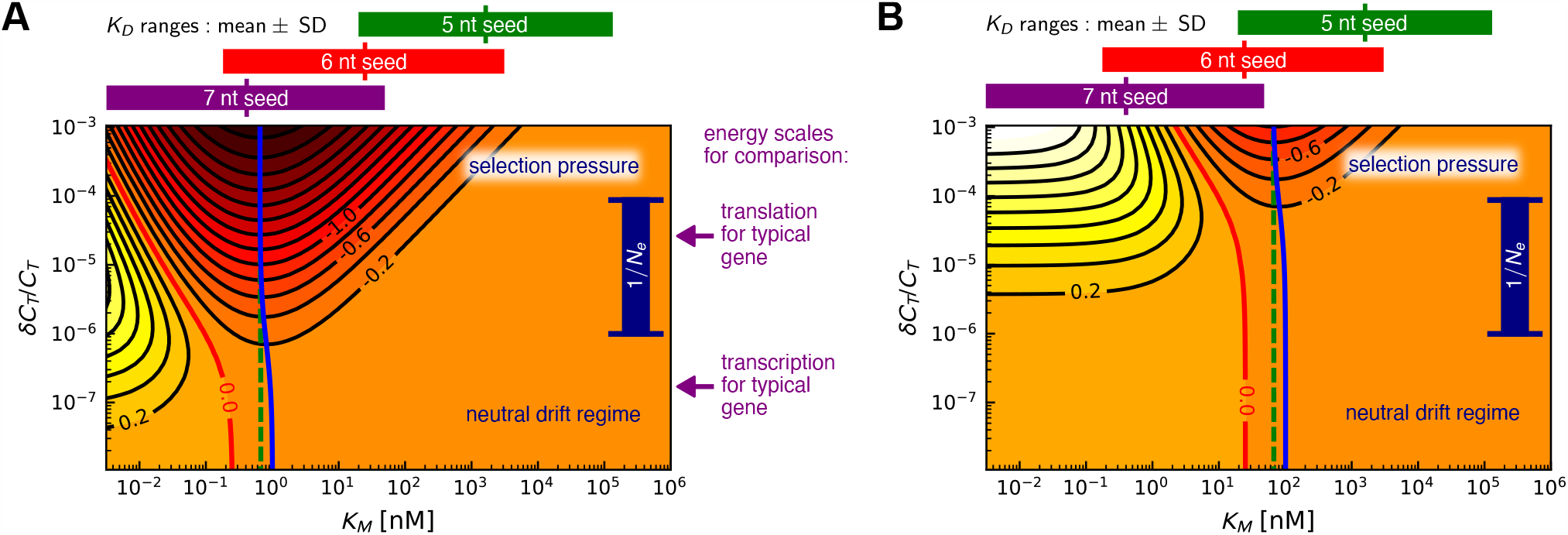
Contour diagrams of noise filter error, log_10_ *E*, in terms of Michaelis-Menten constant *K*_M_ and fractional metabolic cost *δC*_*T*_ */C*_*T*_ for a fixed protein output level 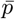 in the (**A**) single target and (**B**) 100 target cases. The spacing between contour values is 0.2. The minimum contour line for a given *δC*_*T*_ */C*_*T*_ corresponds to the most energetically efficient noise reduction, and this is achieved by 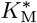 values along the blue curves. Similarly, the green dashed curves show the most efficient 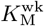 values predicted by the WK optimal filtering theory. The red line is the boundary of the “dud” region on the left, inside which the miRNA regulation adds noise (*E >* 1) rather than mitigating it. To make a connection to physiological values of *K*_M_, we plot estimated *K*_*D*_ ranges for known miRNA-mRNA interactions above the plots for 5, 6, and 7 nt seed lengths. *K*_D_ sets a lower bound on *K*_M_ from Eq. [1]. Altogether, this shows that for a single typical target gene, the most economical noise reduction is likely to occur for 7 nt seeds, while 6 nt seeds become favorable for miRNAs with many targets. To better illustrate the evolutionary relevance, we also plot the dark blue bar showing inverse effective population sizes 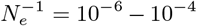 typical for metazoans. As the fitness disadvantage due to metabolic costs become significant, 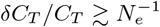, there is selective pressure on the organism driving it towards the 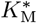 value. For comparison, we also show typical translation and transcription energy scales, based on Ref. [18].

In the biologically relevant parameter regime *γ*_*μ*_, *σ*_*ϵ*_, *ϕ* ≪ 1, where *ϕ* ≡ *d*_*p*_*/d*_*m*_, we can derive analytical approximations for both 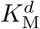 amd 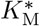. For small costs *δC*_*T*_ */C*_*T*_ we have 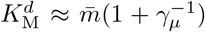, and with increasing cost the boundary begins to decrease as 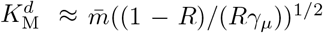. On the other hand, 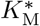 is remarkably stable as *δC*_*T*_ */C*_*T*_ is varied. In the large cost limit it approaches

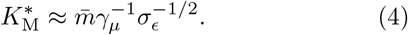

Thus the optimal affinity depends on two non-dimensional biochemical ratios, *γ*_*μ*_ and *σ*_*ϵ*_, and the mean mRNA target number 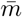.

Fig. 2B shows what happens when we scale up 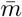 by a factor of 100, roughly mimicking the case of 100 similar mRNA targets. As predicted by Eq. [4], 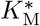 increases by a factor of 100, but the contour levels are pushed up by a similar factor: as expected, it costs more achieve the same level of noise control when compared to the single-target case. Interestingly, the 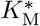 values in the single and multi-target case (about 0.65 nM and 65 nM respectively at high *δC*_*T*_ */C*_*T*_), are comparable to *K*_M_ measured in a fruit fly RNA interference pathway [19]. In the experiments, *K*_M_ = 1 ± 0.2 nM was found for a fully complementary interaction with a 7 nt seed, while single nucleotide mismatches in the seed binding (which in principle allow for a larger range of possible targets) boosted *K*_M_ by up to a factor of 82. While we do not yet have extensive experimental surveys of *K*_M_ values in miRNA (or related siRNA [small interfering RNA]) systems, it would be intriguing to check whether *K*_M_ tends to scale with the target population, as predicted by the optimal theory.

We can make the connection between the number of complementary matches (or seed length) and *K*_M_ more explicit. In SI Sec. S.IV we used the ViennaRNA server [22] to predict the Δ*G*^0^ values of ≈ 10^4^ human miRNA seed sequences of length 7 nt, resulting in a distribution of *K*_D_ values which covered around ten decades on a logarithmic scale. The mean and standard deviation of log_10_(*K*_D_*/*[1*M*]) = −9.5 ± 2.2 is shown as a purple bar above the plots in Fig. 2A,B. To mimic shorter seeds, we deleted 1 or 2 nucleotides from the sequence, to give the ranges log_10_(*K*_D_*/*[1*M*]) = −7.6 ± 2.0 (6 nt, red bar) and −5.8 ± 1.9 (5 nt, green bar). As validation, the calculation was able to correctly reproduce measured *K*_D_ ranges for fully matched 7 nt seeds in fruit fly and mouse siRNA-target complexes. Since *K*_D_ sets the floor for *K*_M_, we see from Fig. 2A that the optimal 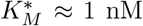 for the single-target case is unlikely to be accessible for 5 nt seeds. It becomes plausible for 6 nt seeds, and even more so for 7 nt seeds. Since *K*_M_ can be up to two decades larger than *K*_D_ [19], it is notable that 7 nt seeds densely cover the range of *K*_D_ values one or two decades smaller than 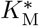. Extrapolating this pattern, longer seeds (≥8 nt) will be less likely than 7 nt to achieve 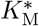. For the 100 target case (Fig. 2B) we see that the ideal seed length is shifted to 6 nt as a result of the increase in 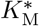. Seeds of length 6 nt constitute 67% of a dataset of human and mouse miRNA seed sequences, with 7 nt seeds forming another 23% [23]. The preponderance of 6 nt seeds is in line with expectations if *K*_M_ was optimized, particularly since miRNAs will typically have many different targets.

## Noise reduction can approach optimal linear filter performance

The filter analogy in Eq. [2] allows us to make an interesting comparison. For a given signal *s*(*t*) and *n*(*t*), we know that there is a filter function *H*_wk_(*t*), the Wiener-Kolmogorov (WK) solution [24–26], which gives the best performance (smallest *E*) of all possible functions *H*(*t*) for this type of linear noise filtering system (SI Sec. S.II B). We denote the corresponding value of error, 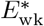, and it serves as the overall lower bound on *E*. In our system, *E* reaches a minimum *E*^*^ at 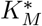 for a given energetic cost, but is this minimum *E*^*^ comparable to 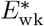? In other words, can miRNA noise regulation approach an optimal WK filter? This type of comparison has recently proven fruitful in a variety of biological contexts [17, 27–33], for example yielding tight bounds on the fidelity of information transmission in signaling networks.

The miRNA system does not exactly realize true WK optimality, because the optimal filter function *H*_wk_(*t*) cannot be precisely implemented by the miRNA regulation network. However, the affinity 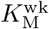 predicted to be optimal by the WK theory for a given *δC*_*T*_ */C*_*T*_ (green dashed curve in Figs. 2A,B) is extremely close to 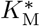 (blue curve). Fig. 3 shows the difference between the respective errors *E*^*^ and 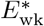 along these curves, as a function of *δC*_*T*_ */C*_*T*_. While 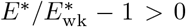, as expected, the difference is always smaller than 0.045, peaking at moderate cost values. As the cost *δC*_*T*_ */C*_*T*_ increases, *E*^*^ converges to 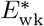, and 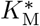 similarly converges to 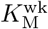. Notably, despite its constraints, the miRNA system can get quite close to WK optimal performance.

**FIG. 3.**
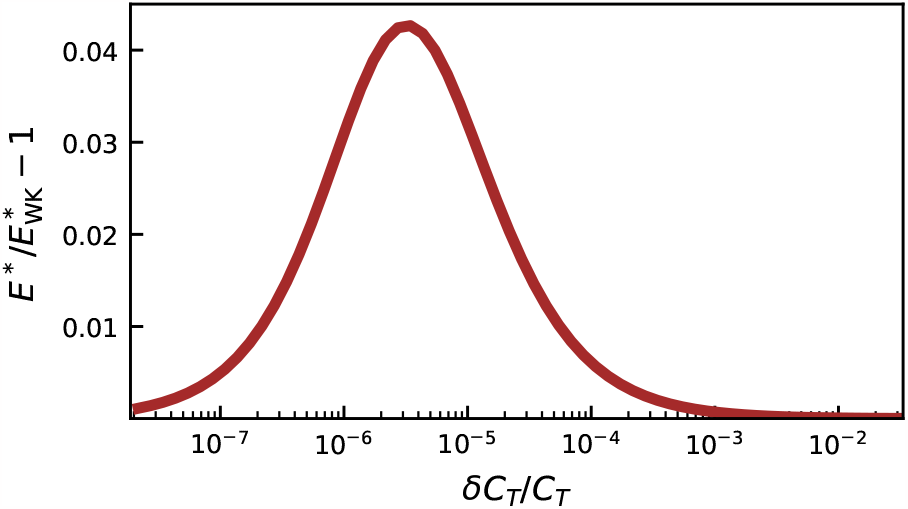
The discrepancy between *E*^*^ and 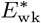, the error of the actual system and the WK optimal system respectively, at the most energetically efficient *K*_M_ for the single target case (Fig. 2A), at different values of the fractional cost 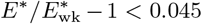, so the miRNA system exhibits a performance close to WK optimality for this network motif.

## Evolutionary pressure on miRNA noise regulation

The previous two sections have argued that optimality in noise reduction (in the broader WK sense) is in principle approachable, and 6-7 nt seeds put miRNA systems within reach of achieving it. The final question we would like to consider is whether there would be any pressure from natural selection actually driving miRNA regulation toward optimality. From a population genetics perspective, let us say that the fitness of an organism with a particular miRNA regulatory network is *f*_*R*_, while the same organism missing the network has fitness *f*_0_. The selection coefficient *s* = *f*_*R*_*/f*_0_ − 1, quantifying the relative fitness, can be decomposed into two contributions, *s* = *s*_*a*_ +*s*_*c*_ [18]. Here *s*_*a*_ is the adaptive advantage due to the regulation, for example resulting from protein noise reduction. The remaining part, *s*_*c*_ *<* 0, comes from the added metabolic cost of implementing the regulation. In order for *s* to be overall positive (and hence *f*_*R*_ *> f*_0_) we would need an *s*_*a*_ *>* 0 that is larger in magnitude than the cost, *s*_*a*_ *>* |*s*_*c*_|.

The evolution of miRNA systems, however, poses a conundrum: a newly arisen miRNA, before any selective fine-tuning of its seed sequence, could potentially target hundreds of mRNA in a random manner, making *s*_*a*_ *<* 0 due to deleterious effects on existing genetic networks. So how would advantageous miRNA regulation eventually emerge? The solution to this problem, as argued by Chen and Rajewsky [2], is for new miRNA systems to be expressed at very low levels, such that *s*_*a*_, even if negative, has a negligible magnitude. In this regime however *s* ≈ *s*_*c*_ *<* 0, which is still an overall fitness disadvantage. Would such a deleterious variant survive in a population long enough for further mutations to confer a significant positive *s*_*a*_? The answer depends on the magnitude of *s*_*c*_ relative to a threshold known as the “drift barrier” [34]. This threshold is set by 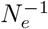, where *N*_*e*_ is a measure of genetic diversity known as the effective population size [35]. *N*_*e*_ is the size of an idealized population that shows the same changes in genetic diversity per generation (due to genetic drift, or random sampling of variants) as the actual population. *N*_*e*_ is generally smaller than the real population size, and among more complex eukaryotes (like vertebrates) can be as small as 10^4^ − 10^6^ [18, 36]. If 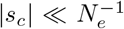, selection against the variant is weak, even when *s*_*c*_ *<* 0, so it can survive in a population via genetic drift as an effectively neutral mutant and even eventually take over (with a fixation probability roughly given by its initial fraction in the population [37]). On the other hand, if 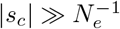, then the costs are sufficiently high that selection would efficiently weed out the variant from the population, unless there were compensating advantages, *s*_*a*_ ≳ |*s*_*c*_|.

A recent derivation based on a general bioenergetic growth model for organisms allows us to make the above discussion more quantitative: it showed that *s*_*c*_ ln(*R*_*b*_)*δC*_*T*_ */C*_*T*_, where *R*_*b*_ is the mean number of offspring per individual [38]. Assuming that ln(*R*_*b*_) does not change the order of magnitude, we can thus use the fractional cost *δC*_*T*_ */C*_*T*_ as a proxy for *s*_*c*_, and compare it to 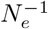. The blue range bars on the right in Fig. 2A,B show possible 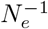 values 10^−6^ − 10^−4^ for higher-order eukaryotes, separating a selection pressure regime at large *δC*_*T*_ */C*_*T*_ from a neutral drift regime at low *δC*_*T*_ */C*_*T*_. For comparison we also show the cost scales for transcription and translation of a typical gene [18], which indicate that transcription is not generally not under selective pressure in these organisms, while translation may be.

At the lowest expression levels, a newly evolved miRNA system could survive in the neutral drift regime, even with a non-optimal *K*_M_. There is limited noise reduction achievable in this regime, since the contours indicating *E* significantly smaller than 1 require larger *δC*_*T*_ */C*_*T*_, particularly for the multi-target case (Fig. 2B). Thus the initial evolution could be imagined as a random walk near the bottom edge of the diagram. Mutations that led to greater expression of an miRNA, moving up on the cost scale, would hit against the drift barrier, and would more likely survive if they came with compensating fitness advantages. In a context where noise control was beneficial, this would mean being funneled up the region where *K*_M_ is close to 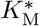, the path that confers the largest noise reduction as expression levels rise. Once the costs are above the drift barrier, there would be significant selective pressure to optimize *K*_M_. In this high cost regime there is a direct tradeoff between extra metabolic expenditure and noise filtering, with Δ*M*_*ν*_ ∼ *M*_*ν*_*/E*, and hence *s*_*c*_ ∼ −*M*_*ν*_/*E*. Our theory predicts that any compensating fitness advantage *s*_*a*_ *>* 0 would have to grow like *E*^−1^ or faster as *E* decreases, in order for the regulation to be viable in the long term.

## Inferring closeness to optimality in experimental systems

Given the achievability of optimal affinities 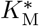 for noise filtering (based on seed lengths), and the evolutionary pressures that could drive a system toward this optimality, is there any corroborating experimental evidence? While we do not have simultaneous measurements yet of *K*_M_ and noise regulation in specific systems, there is a way of approximately inferring the ratio 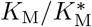 from fitting our model to existing data on miRNA noise suppression. The two experiments we focus on are assays that take the 3’UTR regions of endogenous mRNAs and combine them with fluorescent reporters like *mCherry*—in one case the 3’UTR was from the *Lats2* gene in mouse embryonic stem cells [5], and in the other from the *sens* gene in *Drosophila* wing disc cells [39]. These 3’UTR regions have binding sites for endogenous miRNA, and the assay demonstrates that miRNA interaction suppresses protein noise for these genes, by quantifying reporter protein fluctuations in the wildtype compared to a mutant system where the binding sites were altered, inhibiting miRNA regulation. In both cases the noise suppression has potential functional roles—Lats2 is involved in regulating the cell cycle, apoptosis, and differentiation [40, 41], and excess noise in Sens protein levels leads to disordered sensory patterning in wing disc cells [39].

Thus there is reason to suspect that there could have been selective pressure on the mRNA-miRNA affinity for these genes. To investigate this hypothesis, we took the available experimental data from the two studies and extracted the error *E* and regulation strength *R* as a function of protein expression, since there was a distribution of values for 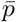 (or reporter intensity) ∝ 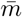 among the population of cells in each system. (The full details of the data analysis can be found in SI Sec. S.V). For a given cell, we know that 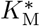 is proportional to the mRNA concentration 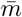 from Eq. [4], and thus should be higher in high-expression versus low-expression cells. On the other hand, the types of miRNA and binding sites are the same between cells, so the actual affinity *K*_M_ should also be the same in each cell. As before, we use a simple version of the multi-miRNA, multi-target theory assuming similar biochemical parameters for each miRNA-mRNA interaction, so we can interpret *K*_M_ to be an average affinity across the ensemble of miRNAs interacting with the targets. To summarize, *K*_M_ is fixed but 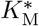 varies between cells, and we can ask whether the ratio 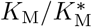 is close to 1 (optimality) over the cell population. Fig. 4 shows the results for this ratio as a function of protein expression in the *Lats2* and *sens* systems, derived from fitting our model to the data. The points connected via lines correspond to fits using the typical parameter values in SI Table S1, while the surrounding colored regions represent uncertainties due to imperfect knowledge of the parameters *γ*_*μ*_ and *σ*_*ϵ*_ (see SI Sec. S.V). Despite this uncertainty, the inferred 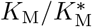 ratio is roughly within an order of magnitude of 1, and for the *Lats2* system even crosses 1. The horizontal dashed lines above and below 1 indicate a factor of 82 in either direction, representing the maximum magnitude of fold change in *K*_M_ observed from a single nucleotide difference in seed matching in another experiment [19]. This range emphasizes the relative narrowness of the observed 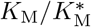 ratio, with the affinity fine-tuned to within a nucleotide of optimality. Moreover, this is true across the whole span of protein expression observed in the experiments, which is also physiologically relevant for these cells. So even though low-expression cells and high-expression cells have different optimal parameters (and error bounds *E*^*^), they all are fairly close to their respective optima. While there is still work to be done in narrowing down parameter uncertainties and extending the analysis to other systems in future studies, this initial analysis is consistent with evolutionary pressures shaping miRNA-mRNA affinity in systems where noise reduction is important.

**FIG. 4.**
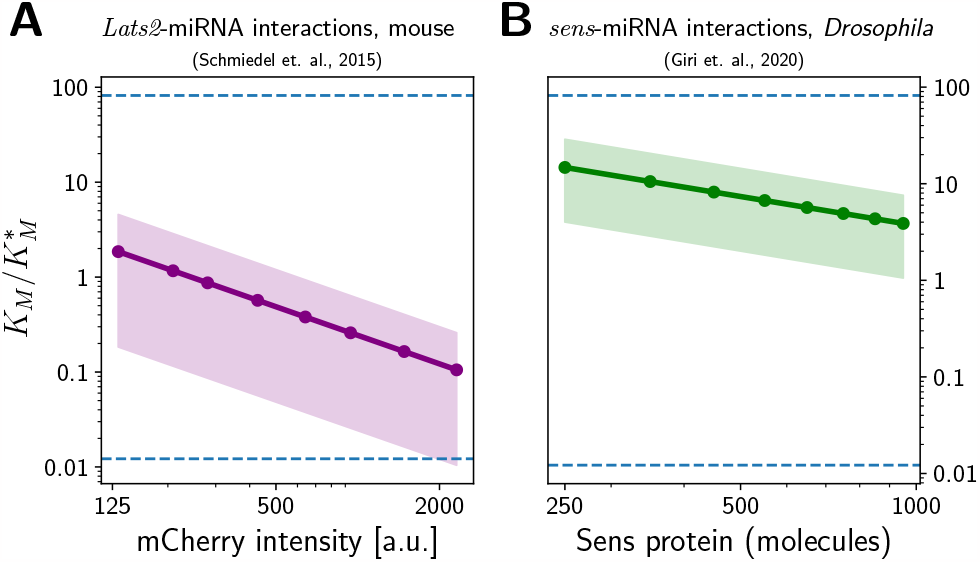
Analysis of optimality for miRNA-mRNA affinity in data from two experimental systems. We show the inferred ratio of actual to optimal Michaelis-Menten constants 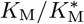, as a function of protein expression level in individual cells, for miRNA regulation of: (A) the *Lats2* gene in mouse embryonic stem cells [5]; (B) the *sens* gene in *Drosophila* wing disc cells [39]. The points connected by a line are theoretical best-fits using typical parameter values from SI Table S1, while the colored regions represent uncertainties due to varying the system parameters *γ*_*μ*_ and *σ*_*μ*_ over biologically plausible ranges. The horizontal axis reflects different quantification of protein expression in the two experiments, either in fluorescent reporter intensity (Lats2) or copy number (Sens). The dashed lines represent the maximum positive or negative fold-change in *K*_M_ observed experimentally from single nucleotide differences in seed matching in Ref. [19].

## Discussion and conclusions

Putting all the elements of our theory together, we show that if noise control confers a fitness advantage for a particular miRNA system, there is selective pressure driving it toward an optimal miRNA-mRNA affinity, as described by the Michaelis-Menten constant *K*_M_. Remarkably, the optimal value 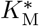 in Eq. [4] can be achieved by a fairly narrow range of seed lengths (6-7 nt), which happen to make up the vast majority of miRNA seeds. While we argue the plausibility of this key result based on realistic ranges of biological parameters, Eq. [4] opens the way for future experimental validation in specific, fully characterized systems. Such validation should be practical, since the equation only involves a small number of biochemical parameters. The noise reduction at optimality approaches the performance of a Wiener-Kolmogorov filter, the best possible linear noise filter. If true, such optimization would be a striking example of metabolic costs directly shaping the course of evolution for a biochemical network in eukaryotes. This is unusual in itself, because eukaryotes are generally less likely to prioritize energy efficiency relative to prokaryotes, which have higher effective population sizes and thus lower drift barriers [18]. Beyond the implications for miRNA evolution, the theory could also find applications in the design of synthetic circuits with 3’UTR engineering and artificial miRNAs [42–45].

While the simple theory of miRNA-mRNA interaction used here is sufficient to describe certain experiments [5], there are a variety of model assumptions that can be relaxed in future investigations, to test the conclusions more broadly. For example, one aspect was ignored in the current model: many miRNAs can bind to multiple sites on a single target, with potentially different affinities, as well as exhibit varied affinities more generally across multiple mRNA targets. While we do not expect this heterogeneity to qualitatively change the overall results, it will be interesting to see how it shifts the relations between affinity, metabolic costs, and noise reduction. Multiple binding sites / targets can also lead to nonlinear phenomena like ultrasensitivity and bistability in miRNA-mRNA systems [46], which in turn could give noise reduction additional functional implications, like inhibiting stochastic switching between different cellular states. More complex models can also consider the role of miRNA in mitigating the effects of (possibly nonstationary) environmental fluctuations, along with the noise due to cellular processes [8, 47].

Though our work focuses on noise control via miRNA regulation, it is also important to keep in mind that noise control can be implemented via other mechanisms, and that miRNA themselves have other functional roles. Protein noise can be reduced (maintaining the same expression level) by increasing transcription and either decreasing translation rates or enhancing mRNA degradation [48, 49], and this would avoid noise added due to miRNA interactions [50]. Degradation rates could for example be fine-tuned by the length of mRNA poly(A) tails [51]. However this mechanism is non-specific, while miRNAs allow for selective noise control via seed recognition. Though our results suggest a formative role for noise reduction in shaping miRNA-mRNA affinity, the filtering capacity likely co-evolved with other miRNA functions such gene silencing and crosstalk [52]. Perhaps in some cases noise regulation is simply a complementary benefit of a far more complex utility scheme.

Take for instance the scenario where there is selective pressure for gene silencing via miRNA, in other words suppressing the protein level 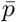 for a particular gene. Naively, one might expect this pressure to always favor smaller values of *K*_M_, since higher affinities can provide the same amount of suppression using fewer miRNA (and hence at smaller metabolic cost). But as we saw in our theoretical analysis (i.e. the “dud” regime on the left in the panels of Fig. 2), smaller *K*_M_ also introduces extra noise into the system. If this noise becomes sufficiently high that it has deleterious effects, like unwanted stochastic transitions between cellular states, then there will be a countervailing pressure to increase *K*_M_. The balance between these two effects might result in a metabolic sweet spot for affinity, analogous to the one we described in our model, though the posited functional role of the miRNA system was different.

Viral miRNAs present another example of alternative functional roles. These miRNAs exploit the host metabolism, and are likely useful for non-noise-related tasks like evading the host immune response [53]. Notably, though viruses lack their own metabolic machinery, there can still be selective pressures on viral miRNA expression as part of the overall energetic costs associated with viral copying [54]. Ultimately a deeper understanding of miRNA evolution will require larger-scale models of its full regulatory context, coupled with *in vivo* experiments to explore the tangled effects of function, metabolic costs, and fitness.

## ACKNOWLEDGMENTS

We thank Amit Singh Vishen for helpful discussions and Xingbo Yang for a critical reading of the manuscript and insightful comments.

## Data availability

Code related to the calculations, routines for plotting the figures, and all the associated datasets are available at the following Github repository: https://github.com/hincz-lab/microrna/.

## Supplemental Material

### S.I ANALYSIS OF MICRORNA-MEDIATED NOISE REGULATION

#### A Chemical Langevin formalism of microRNA-mediated gene expression

##### 1 General birth-death processes

We start with a general description of chemical Langevin (CL) dynamics, which is the basis for our theoretical model. A biochemical reaction network consists of *N* different types of biomolecules with time-dependent populations *n*_*j*_(*t*), *j* = 1, …, *N*. The rate of change of a given population, *dn*_*j*_(*t*)*/dt*, can be expressed in terms of a number of possible stochastic reactions, which we will label with an index *ρ* = 1, …, *r*, where *r* is the total number of reactions. The CL dynamics of the birth-death process for biomolecule *j* can be written as [1]:

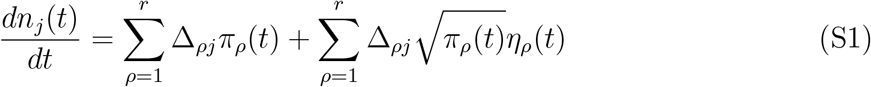

where Δ_*ρj*_ are the stoichiometric coefficients representing the change in the population of *j* in reaction *ρ, π*_*ρ*_(*t*) are the rates of reactions and *η*_*α*_(*t*) are independent Gaussian noise terms at time *t* which satisfy ⟨*η*_*ρ*_(*t*)*η*_*β*_(*t*′)⟩ = *δ*_*ρβ*_*δ*(*t* − *t*′) where *δ*_*ρβ*_ and *δ*(*t* − *t*′) are Kronecker delta and Dirac delta functions, respectively. As the reactions for the production and degradation of *j* are uncorrelated, we can add up the uncorrelated noise terms, and obtain:

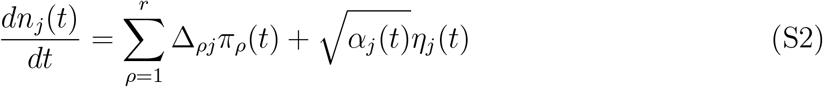

where 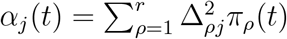. Note that though the *η*_*j*_’s are uncorrelated at different times, there can be non-zero correlations between the dynamics of different populations at time *t* if they share the same reactions.

##### 2 Protein production

To apply the CL framework to microRNA regulation of gene expression, we focus on the following biomolecular species and reactions, following the model of Ref. [2]:

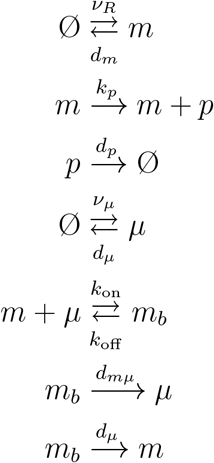

Here *m, p, μ, m*_*b*_ represent respectively mRNA, protein, miRNA and the miRNA-mRNA complex. These reactions include birth-death processes of individual species as well as the formation and degradation of complexes. When the miRNA-mRNA complex *m*_*b*_ degrades, the miRNAs are recycled. For convenience, we assume a similar property when miRNAs spontaneously degrade. This model has been shown to agree well with experimental systems of miRNA-mediated gene expression and noise reduction [2, 3].

The reactions above give the following mass-action kinetics for the concentration3s of the four species (denoted by brackets):

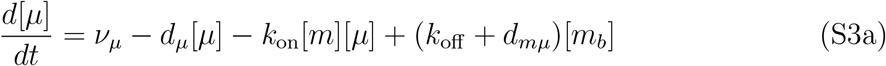

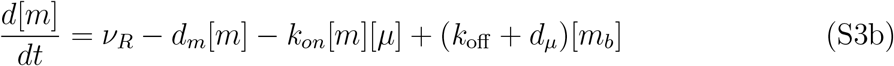

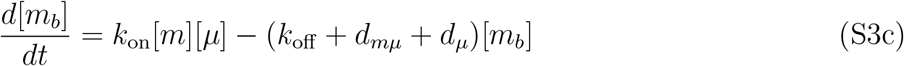

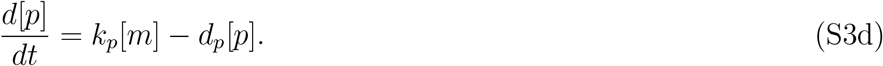

The total miRNA concentration is a sum of free and bound miRNAs, i.e., [*μ*_*tot*_] = [*μ*] +[*m*_*b*_]. Assuming the binding/unbinding rate of the mRNA-microRNA complex is much faster than other dynamics, we obtain a quasi-steady-state approximation leading to 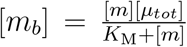, with Michaelis-Menten constant *K*_M_ = *K*_D_ + *k*_cat_*/k*_on_ expressed in terms of the dissociation constant *K*_D_ = *k*_off_ */k*_on_ and an effective catalytic rate constant *k*_cat_ = *d*_*mμ*_ + *d*_*μ*_ for the miRNA-catalyzed degradation reaction. With these simplifications, we obtain

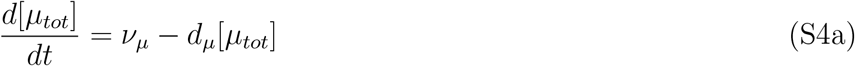

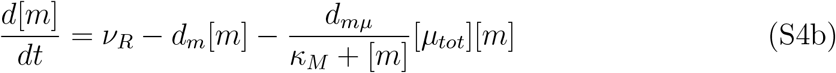

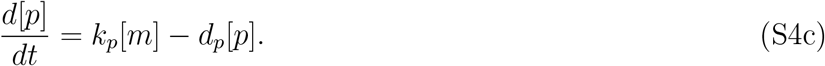

Note that the description in terms of chemical concentrations ignores the fact that the crowded cell interior is not an ideal solution, and that in principle the model can be made more accurate by replacing concentrations with chemical activities in all the reaction rate terms. However, the activity coefficients depend on the concentrations of all chemical species in the cytoplasm (including species not explicitly present in the model), which makes mathematical analysis intractable. The simplest approximation is to assume constant activity coefficients, based on the steady-state values of all the concentrations. In this scenario the activity coefficients can be absorbed into effective rate constants, and the structure of the kinetic equations above stays the same.

Converting concentrations to copy numbers (denoted without brackets) by multiplying with the cell volume *V*, and using Eqs. (S1)-(S2), we can now introduce the corresponding CL equations for the model:

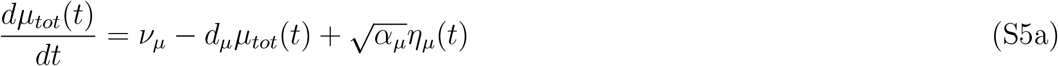

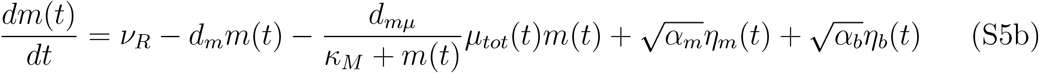

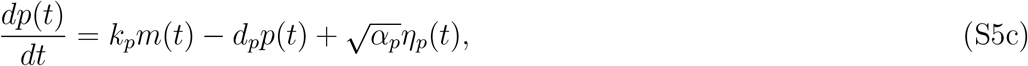

where ⟨*η*_*i*_(*t*)*η*_*j*_(*t*′)⟩ = *δ*_*ij*_*δ*(*t*−*t*′). We will provide a detailed derivation of the noise amplitudes *α*_*j*_(*t*) in Section S.I B.

##### 3 Steady-state properties, regulation strength, bound fraction of miRNAs

At the steady-state, the mean populations for our dynamical system are 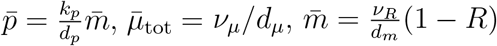, where the regulation strength *R* is defined as:

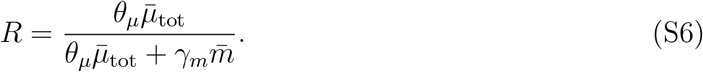

Here *γ*_*m*_ ≡ *d*_*m*_*/d*_*mμ*_ and *θ*_*μ*_ is the bound fraction of miRNAs,

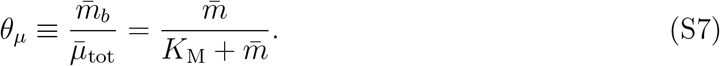

In order to compare protein noise at different levels of miRNA regulation (including the no regulation case), we fix the mean protein population by assuming the mRNA production rate *ν*_*R*_ changes with *R* to compensate for the effects of regulation,

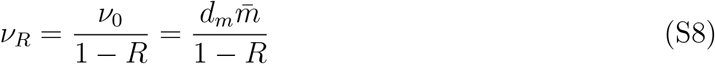

where *ν*_0_ = *ν*_*R*_(*R* = 0)

#### B Protein expression noise

The fluctuating molecular populations are defined in the main text as 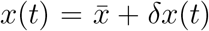 where *δx*(*t*) is the variation from the mean level 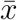 and the species of interest are *x*(*t*) ≡ {*μ*_*tot*_(*t*), *m*(*t*), *p*(*t*)}. Since we focus on the stationary-state fluctuations, we may treat the noise amplitudes *α*_*j*_ as time-independent. We can write these in terms of the equilibrium populations,

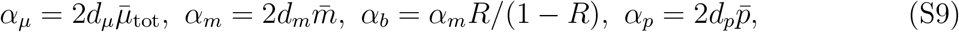

where 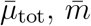, and 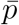 are mean steady-state population levels for miRNA, mRNA and protein respectively. Using the linear-noise approximation in Eq. (S5) and taking the Fourier transform leads to

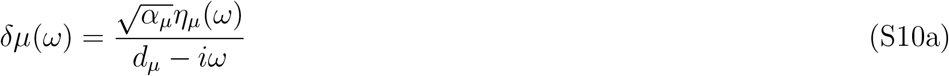

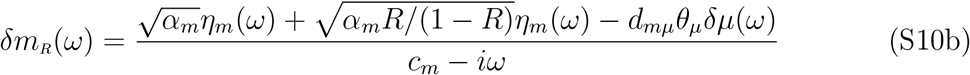

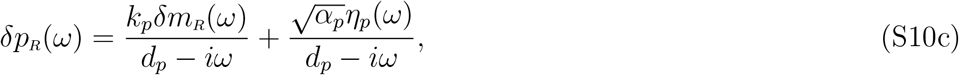

with *c*_*m*_ = *d*_*m*_(1−*Rθ*_*μ*_)*/*(1−*R*). In Fourier space, the noise correlations satisfy ⟨*η*_*i*_(*ω*)*η*_*j*_(*ω*′)⟩ = 2*πδ*_*ij*_*δ*(*ω* + *ω*′). The limit *R* → 0 corresponds to an unregulated system, with *c*_*m*_ → *d*_*m*_ and *d*_*mμ*_*θ*_*μ*_*δμ*(*ω*) → 0 in Eq. (S10b).

The main measure that we use to assess noise levels in protein populations is the Fano factor 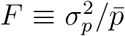 where the variance 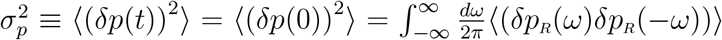. Using Eq. (S10c) and the correlation properties of the noise terms *η*_*i*_(*ω*) in Fourier space, we can calculate the Fano factors *F*_0_ and *F*_*R*_ for the unregulated and regulated systems respectively:

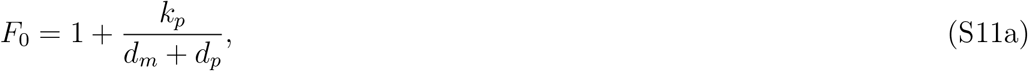

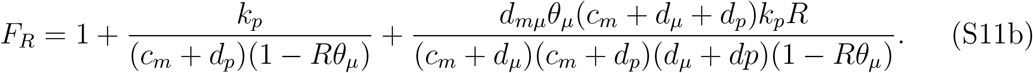

The third term in *F*_*R*_ is a result of coupling with microRNA pool noise (extrinsic noise). We can quantify the relative protein noise levels in an miRNA-regulated system versus an unregulated one with the quantity

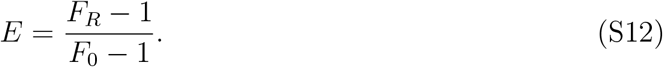

To simplify our analysis, we define the following nondimensional parameters:

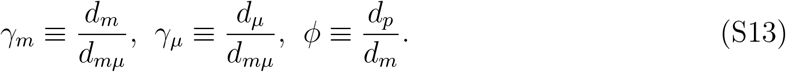

Assuming the degradation rate constant of proteins is slower than that of RNAs, i.e. *ϕ* ≪ 1, we can derive an approximate expression for *E*,

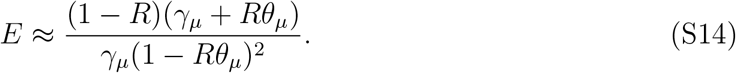

#### C Bioenergetic costs of miRNA regulation

To estimate bioenergetic costs, we follow the Lynch & Marinov analysis [4], where the transcriptional cost of a typical gene in eukaryotic cells is given by *C*_*T*_ ≡ *c*_*g*_ + *M*_*ν*_*t*_*r*_. Here *c*_*g*_ are costs due to synthesis (growth), *M*_*ν*_ is the energy consumption rate for maintenance, and *t*_*r*_ is the lifetime of the cell. For long enough cell-division times, the maintenance term dominates, and hence *C*_*T*_ ≈ *M*_*ν*_. For the unregulated system, this amounts to *M*_*ν*_ = *ν*_0_*ϵ*_*m*_, where *ϵ*_*m*_ is the maintenance energy cost per mRNA (i.e. the turnover cost of recycling its nucleotides after degradation into new mRNA). By contrast, the cost for the miRNA regulated system consists of two contributions: increased transcription of mRNAs in order to maintain 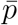 and transcription of miRNAs. The additional cost relative to the unregulated case, Δ*M*_*ν*_, can be expressed as Δ*M*_*ν*_ = (*ν*_*R*_ − *ν*_0_)*ϵ*_*m*_ + *ν*_*μ*_*ϵ*_*μ*_, where *ϵ*_*μ*_ is the maintenance cost per miRNA. By substituting *ν*_*R*_ = *ν*_0_*/*(1 − *R*), 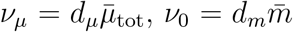 from the definitions in Section S.I A 3, and inverting Eq. (S6) for 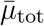, we obtain Eq. (3) in the main text,

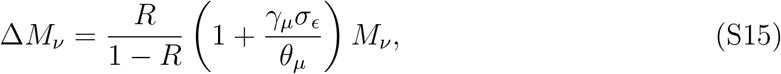

with *σ*_*ϵ*_ ≡ *ϵ*_*μ*_*/ϵ*_*m*_.

In order to determine the noise levels for a given cost Δ*M*_*ν*_, we can invert the above equation for *R*, and then plug into Eq. (S14). In the limit *ϕ* ≪ 1 we get

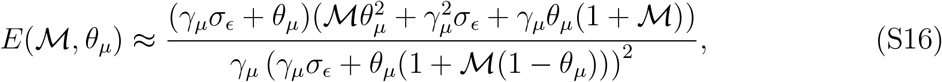

where ℳ = Δ*M*_*ν*_*/M*_*ν*_.

### S.II. THE NOISE-FILTER FORMULATION OF THE MICRORNA REGULATED SYSTEM

#### A Realizing miRNA regulation as a noise filtering mechanism

MicroRNA regulation acts directly on mRNA-related fluctuations in protein output production. We can formulate noise regulation as a filtering process by decomposing the protein fluctuations given in Eq. (S10c) as

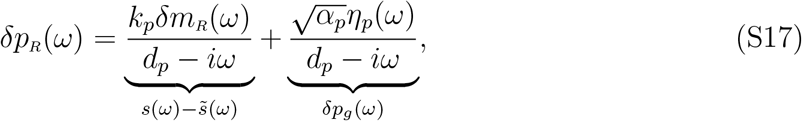

where *δp*_*g*_(*ω*) is the ground noise in protein production (the intrinsic noise independent of mRNA fluctuations), and 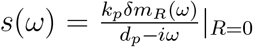 is the contribution from mRNA noise in the absence of regulation. The perturbation due to regulation, 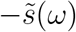, can then be interpreted in the noise filter framework as an “estimate” of the signal *s*(*ω*), with maximum reduction of the mRNA-induced noise occuring when the estimate is perfect, 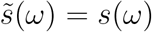. Note that 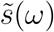 and *s*(*ω*) have non-zero correlations, while 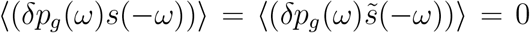. Hence miRNA regulation can compensate for the mRNA noise, but not the ground noise. The normalized error in estimation is defined in the time domain,

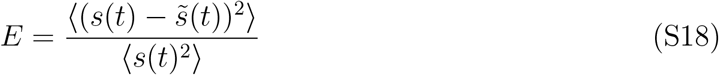

which is equivalent to Eq. (S12) since 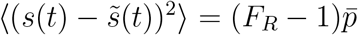 and 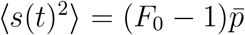. For a general linear filtering problem we can write the estimated signal as

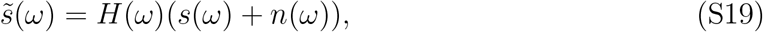

where *s*(*ω*)+*n*(*ω*) ≡ *c*(*ω*) is the signal corrupted with noise *n*(*ω*), and *H*(*ω*) is the linear filter function. Eq. (2) of the main text is the inverse Fourier transform of Eq. (S19), showing filtering as a convolution in the time domain,

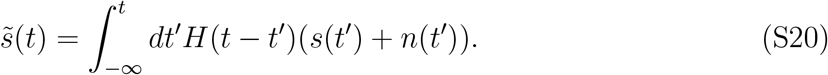

The fact that the upper limit of the integral is *t* reflects an additional physical constraint on the filter: in the time domain *H*(*t*) = 0 for all *t <* 0. This enforces causality, namely that the estimate 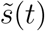 can only depend on on the past history (*t*′ *< t*) of the noise-corrupted signal *s*(*t*′) + *n*(*t*′). In Fourier space, this translates to the requirement that *H*(*ω*) extended to complex *ω* can have no poles or zeros in the upper half-plane, Im *ω >* 0.

For our system, each of terms in Eq. (S19) can be identified using Eqs. (S10b), (S17), and (S19):

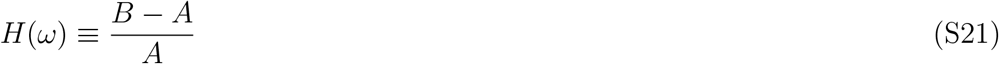

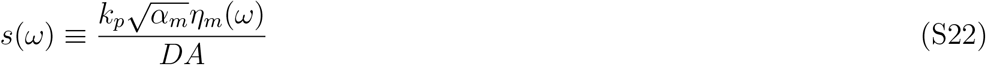

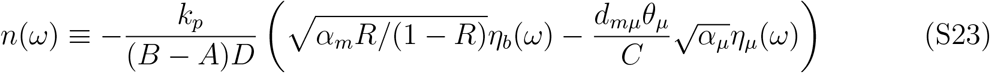

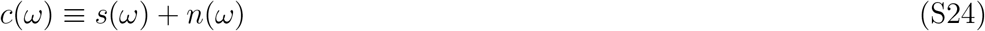

with complex functions

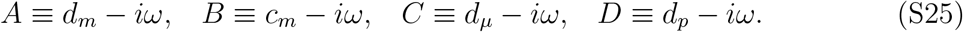

The noise term *n*(*ω*) captures the additional fluctuations due to miRNA-mRNA interactions and it is uncorrelated with the signal *s*(*ω*).

#### B Wiener-Kolmogorov optimality of protein output noise level

Wiener-Kolmogorov (WK) theory provides a framework to calculate the optimal causal filter function *H*_wk_(*ω*) that minimizes the error *E*, and thus represents the fundamental limit on how much linearized regulation can reduce mRNA noise. An overview of the WK formalism can be found in Ref. [5]. Here we show how to apply it to the miRNA system.

In order to determine the error function *E* in Eq. (S18), we need to calculate the variance given in the numerator:

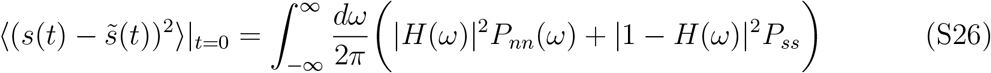

where the power spectra (or cross-spectral density) functions are defined as:

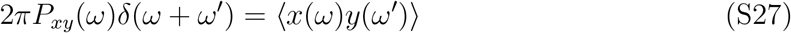

for *x, y* = *s* or *n*. The inverse Fourier transform of this expression would give the cross correlations in the time domain, *C*_*xy*_(*t*) ≡ ⟨*x*(*t*′)*y*(*t*′ + *t*)⟩. The WK optimal filter minimizing *E* for a given *s*(*t*), i.e., minimizing Eq. (S26), satisfies

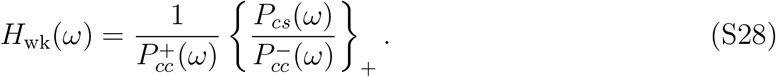

The above expression requires two types of decomposition into causal (+) and anti-causal (−) parts, where a causal term in Fourier space has no poles or zeros in the upper complex *ω* half-plane, and an anti-causual term is the complex conjugate of a causal function. The first decomposition is multiplicative, where we write 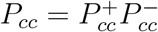, a product of causal and anti-causal terms. The second decomposition is additive, denoted with brackets as 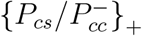. Here {*G*(*ω*)}_+_ for a function *G*(*ω*) can be calculated by doing a partial fraction expansion of *G*(*ω*) can keeping only terms with no poles in the upper half-plane.

The relevant power spectra are obtained by inserting Eqs. (S22), (S23), (S24) into Eq. (S27):

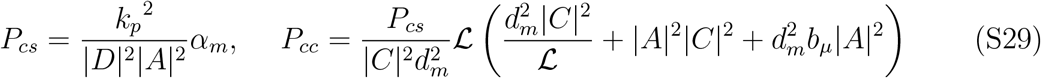

and *P*_*ss*_ = *P*_*cs*_, *P*_*nn*_ = *P*_*cc*_ − *P*_*cs*_ with dimensionless parameters 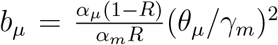 and 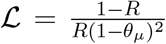. To achieve the causal decompositions required for WK equation (S28), *P*_*cc*_ can be rewritten as,

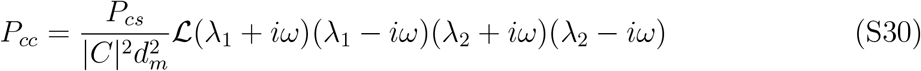

where ±*iλ*_1,2_ are the roots of the numerator and given by

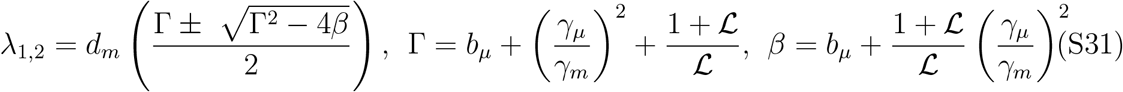

Thus, we obtain

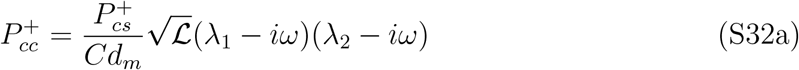

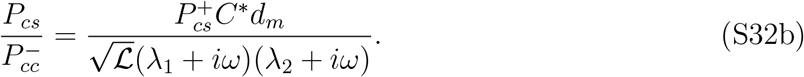

where 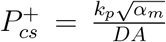. Next, we need to determine the additive decomposition of 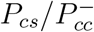, yielding the casual part

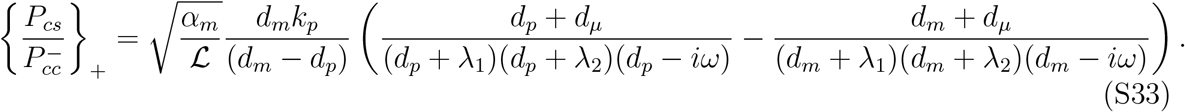

The optimal WK filter can be calculated from Eq. (S28) by inserting Eqs. (S32a) and (S33), resulting in

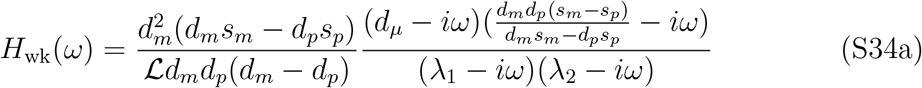

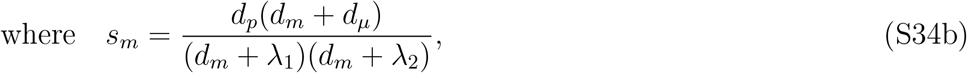

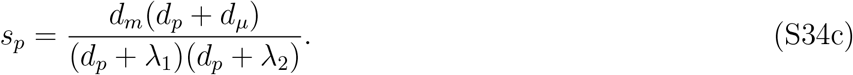

The filter *H*_wk_(*ω*) contains two characteristic frequencies *λ*_1_, *λ*_2_. This differs from the functional form of *H*(*ω*) in Eq. (S21) which has a single characteristic frequency 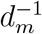. To evaluate how close the miRNA-mediated noise regulation can approach WK optimality, we numerically compare the errors *E, E*_wk_ of both cases respectively. We get the error in estimation in WK optimality using Eq. (S26) in Eq. (S18) with *H*(*ω*) → *H*_wk_(*ω*),

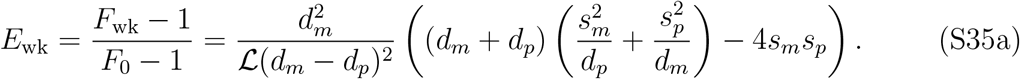

Fig. S1 illustrates the discrepancy between *E*_wk_ from the above equation and *E* from Eq. (S12).

**FIG. S1.**
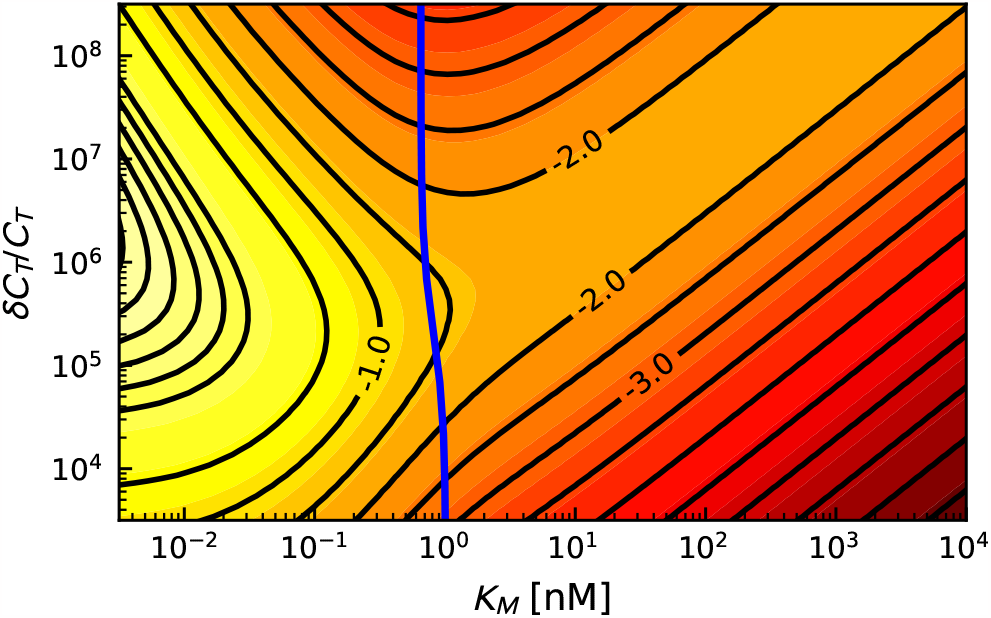
Contour diagrams of log_10_ (*E/E*_wk_ − 1), in terms of Michaelis-Menten constant *K*_M_ and fractional metabolic cost *δC*_*T*_ */C*_*T*_ for a fixed protein output level 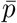 in the single target case using the parameters from Table S1. The contour spacing is 0.5, and the blue line follows the most energetically efficient noise reducing 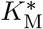 as in Fig. 2 of the main text.

### S.III. CALCULATION OF PHASE DIAGRAMS

#### A Contour plots, fitness costs, and Michaelis-Menten constant

In the main text, we plot the contour diagram of log_10_ *E* from Eq. (S12), describing noise reduction for miRNA-regulated gene expression at a fixed protein output 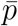. The gene expression parameters are given in Table S1, while the Michaelis-Menten constant *K*_M_ and miRNA levels *μ*_tot_ are taken as free parameters which control both noise reduction and the metabolic costs of regulation. The regulation costs are calculated using Eq. (S15), and the definitions of energetic costs per mRNA and per miRNA respectively,

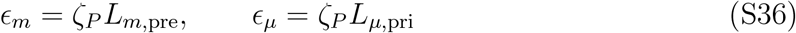

where *ζ*_*P*_ is the energetic need per nucleotide (which we express in units of phosphate bonds hydrolyzed), and *L*_*m*,pre_, *L*_*μ*,pri_ are the numbers of nucleotides of precursor mRNA and primary miRNA. In principle, the costs associated with the miRISC can be larger than *ϵ*_*μ*_. Thus, our estimations can be seen as a lower bound. Finally, we relate metabolic costs to a fitness disadvantage, *s*_*c*_ ∼ Δ*M*_*ν*_*/M*_tot_ where *M*_tot_ is the (mean) total metabolic rate of a cell, as discussed in the main text.

**TABLE S1.** Gene expression parameters (metazoan) used in calculating the phase diagrams. Typical (or estimated) values from the corresponding references are taken such as median values over a distribution of genes. When an extensive distribution is not present, a reasonable estimation is made compatible with the references. We have not found a value for *d*_*mμ*_ in the literature, however, target mRNAs degrade faster in the complex, and hence we assumed the ratio to be within an order of magnitude. In order to convert the copy number units to molar units of mRNA numbers and *K*_m_, we divide these values by the cell volume *V* and Avagadro constant.

| Parameter | Definition | Value | Reference |
| --- | --- | --- | --- |
| $d_m$ | mRNA degradation rate constant | 0.11 hr<sup>-1</sup> per mRNA | [6] |
| $d_p$ | protein degradation rate constant | 0.022 hr<sup>-1</sup> per protein | [6] |
| $d_\mu$ | microRNA degradation rate constant | 0.05 hr<sup>-1</sup> per miRNA | [7, 8] |
| $d_{m\mu}$ | mRNA degradation through miRISC | 0.44 hr<sup>-1</sup> per complex | |
| $K_M$ | Michaelis-Menten const. for target degradation | solved for | |
| $L_{m,\text{pre}}$ | precursor mRNA nucleotide number | 20,000 | [9] |
| $L_{\mu,\text{pri}}$ | primary miRNA nucleotide number | 2,500 | [10, 11] |
| $\zeta_P$ | P bonds hydrolyzed per nucleotide recycling | 2.17 P hr<sup>-1</sup> | [4] |
| $M_{\text{tot}}$ | total metabolic rate per cell | $3 \times 10^{11}$ P hr<sup>-1</sup> | [4] |
| $V$ | volume of cells | 2,000 $\mu\text{m}^3$ | [12] |
| $\phi$ | $d_p/d_m$ | 0.2 | |
| $\gamma_m$ | $d_m/d_{m\mu}$ | 0.25 | |
| $\gamma_\mu$ | $d_\mu/d_{m\mu}$ | 0.114 | |
| $\bar{m}$ | mean mRNA copy number | 31 | [6, 12] |
| $\bar{p}$ | mean protein copy number | 200,000 | [6, 12] |
| $\sigma_\epsilon$ | $\epsilon_\mu/\epsilon_m$ | 0.125 | |

#### B Minimal noise for the same metabolic cost

The contour plots indicate a minimum *E* for a given metabolic cost ∼ Δ*M*_*ν*_ at a precise *K*_M_ value. This value remains in a narrow range as a function of metabolic costs as shown in the Fig. 2 of the main text. Taking the limit *ϕ* ≪ 1, we can estimate this value by minimizing Eq. (S16) with respect to *θ*_*μ*_(*K*_M_). Among multiple solutions, there is a single physical one, from which we can infer 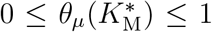 i.e., 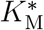 using Eq. (S7). For high ℳ and *γ*_*μ*_ ≪ 1, this yields the Eq. 4 of the main text,

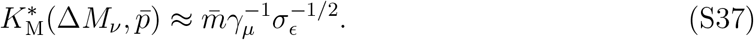

This approximation holds well for the parameters in Table S1, with only 0.6% discrepancy at high ℳ. For *σ*_*ϵ*_ ≪ 1 we observe that 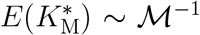, which relates noise reduction to metabolic and hence to the fitness costs, i.e., 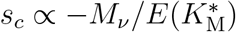.

### S.IV. ESTIMATING DISSOCIATION CONSTANTS *K*_D_ FOR SEED SEQUENCES OF VARYING LENGTH

To estimate a range of possible *K*_D_ values for miRNAs with different seed sequences, we started with a collection of 9990 human miRNA seed sequences of length 7 nucleotides (nt) taken from the TargetScanHuman database, ver. 7.1 [14]. For each sequence, we used the RNAcofold algorithm from the ViennaRNA package (ver. 2.3) to calculate the free energy Δ*G*^0^ at *T* = 298^*°*^K for binding to a complementary target sequence [13]. To mimic the effects of unpaired nucleotides flanking the seed-target complex, we added two random unpaired nucleotides at both the beginning and end of the seed/target to make the total length 11 nt. The algorithm was run with the default dangling end energy option. Once Δ*G*^0^ is known, *K*_D_ can be found using the relation Δ*G*^0^ = *k*_*B*_*T* ln(*K*_D_*/*[1*M*]).

To validate the predictions of the algorithm, we checked it on the seed sequence GAG-GUAG, for which experimental *K*_D_ values are available for fruit fly and mouse siRNA-target complexes [15]. Depending on the dangling ends, the algorithm predicted a range of *K*_D_ with mean 63 pM (95% confidence interval: 7 to 238 pM). This compares well with the experimentally measured ranges for different seed-matched targets: 4 to 210 pM for fly, 13 to 26 pM for mouse [15].

A histogram of the results for all the seed sequences in the dataset is shown in Fig. S2A. The distribution has 8 peaks, roughly corresponding to the fact that the seed-target complex can have between 0 and 7 CG pairs. The more CG pairs relative to AU pairs, the stronger the binding and the lower the *K*_D_. To model the effect of having a 6 nt seed, in Fig. S2B the calculation was redone after randomly deleting 1 nucleotide from each original seed. Similarly Fig. S2C shows the results for 5 nt seeds, after randomly deleting 2 nucleotides per seed. As expected, shorter seeds lead to overall weaker binding, shifting the distribution to larger *K*_D_ values.

### S.V. ANALYSIS OF EXPERIMENTAL MICRORNA SYSTEMS

In this section we use our theoretical framework to analyze data from two earlier experimental works on microRNA-mediated noise regulation. As mentioned in Sec. S.I., the basic mathematical framework for miRNA-regulated protein production, which is the starting point for our theory, has already been validated in Ref. [2] in a variety of experimental scenarios. Here we show another aspect of the analysis, interpreting the experimental data via the optimal noise filter formulation of the theory. We focus on two systems, both involving endogenous miRNAs in living cells interacting with binding sites on 3’UTR regions from endogenous genes, with the latter coupled to fluorescent reporters that allow direct measurements of regulation strength and noise reduction. The genes are: i) *Lats2* in mouse embryonic stem cells (mESCs) [2]; ii) *sens* in *Drosophila* wing disc cells [16].

**FIG. S2.**
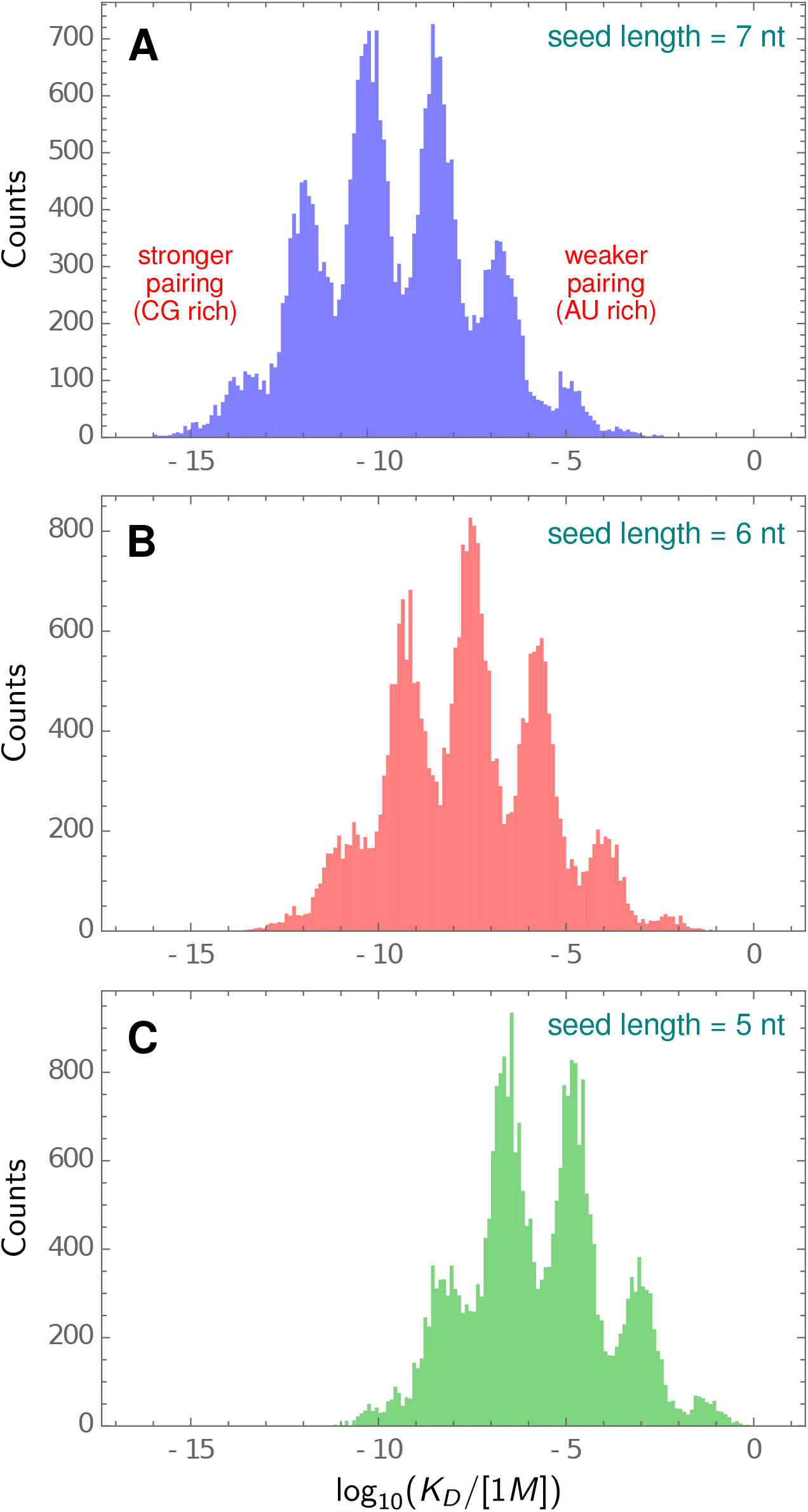
Distribution of *K*_D_ values estimated for a different miRNA seed sequences with length: A) 7 nt; B) 6 nt; C) 5 nt. The calculation used the RNAcofold algorithm of the ViennaRNA package [13].

Let us start with an overview of the analysis procedure, before turning to specific details for the two systems. In each experiment there is a comparison between a miRNA-regulated system, and one without regulation, where the 3’UTR target sites are modified to prevent miRNA binding. This allows us to extract the error *E* from the relative noise levels in the regulated versus unregulated case, as defined in Eq. (S12), as well as the regulation strength *R*. The experimental measurements are carried out over a population of cells where the protein expression levels (as quantified by the fluorescent reporter) vary, and we will use *X* to denote the measure of protein levels in a cell, with *X* corresponding to fluorescence intensity (in arbitrary units) in the *Lats2* case, and to protein numbers in the *sens* case (where the calibration between intensity and copy numbers was carried out). The noise filter characteristics in a given cell will depend on 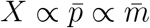, and hence the experimental values of *E*^exp^(*X*) and *R*^exp^(*X*) will both vary with *X*. In our analysis, we fit *E*^exp^(*X*) to Eq. (S14), our analytical result in the limit where the protein degradation rate constant is much smaller than the mRNA degradation rate coefficient (*ϕ* ≪ 1). Plugging Eq. (S7) and *R*^exp^(*X*) into Eq. (S14), we get the following theoretical expression relating the two measured quantities:

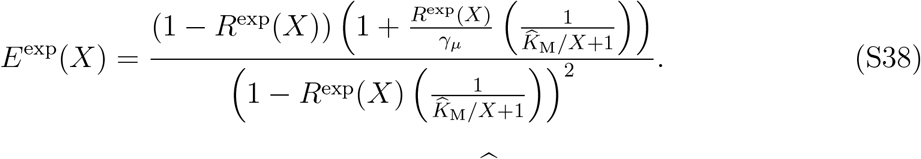

where we define a rescaled Michaelis-Menten constant 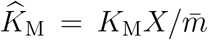, with 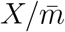 being the constant scaling factor between *X* and mRNA number 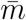. Eq. (S38) has only two unknowns, *γ*_*μ*_ and 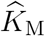, which can be estimated via fitting to the experimental data for *E*^exp^(*X*) and *R*^exp^(*X*) as a function of *X*. Once 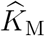 is determined from the fit, we can use main text Eq. [4] for 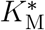 to find the ratio of actual to optimal Michaelis-Menten constants,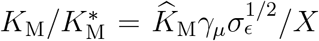, which is plotted in main text Fig. 4. Note that this ratio scales inversely with expression level, 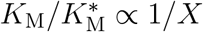, and thus cells with different *X* will be closer or further away from optimality. As discussed in the main text, the results demonstrate that across a physiological range of protein expression levels, both experimental systems exhibit *K*_M_ roughly within an order of magnitude of 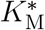.

#### miRNA regulation of *Lats2* in mouse embryonic stem cells [2]

The regulation strength *R* is obtained by the relative expression levels in regulated and unregulated cases, i.e., 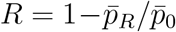. This was determined via a dual reporter system and is almost constant at *R*^exp^(*X*) ≈ 0.8 for all expression levels *X* (Fig. S3A), quantified in terms of the fluorescent marker (mCherry) intensity *X*. The protein expression noise in these experiments was reported in terms of the coefficient of variation,

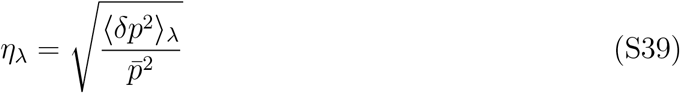

**FIG. S3.**
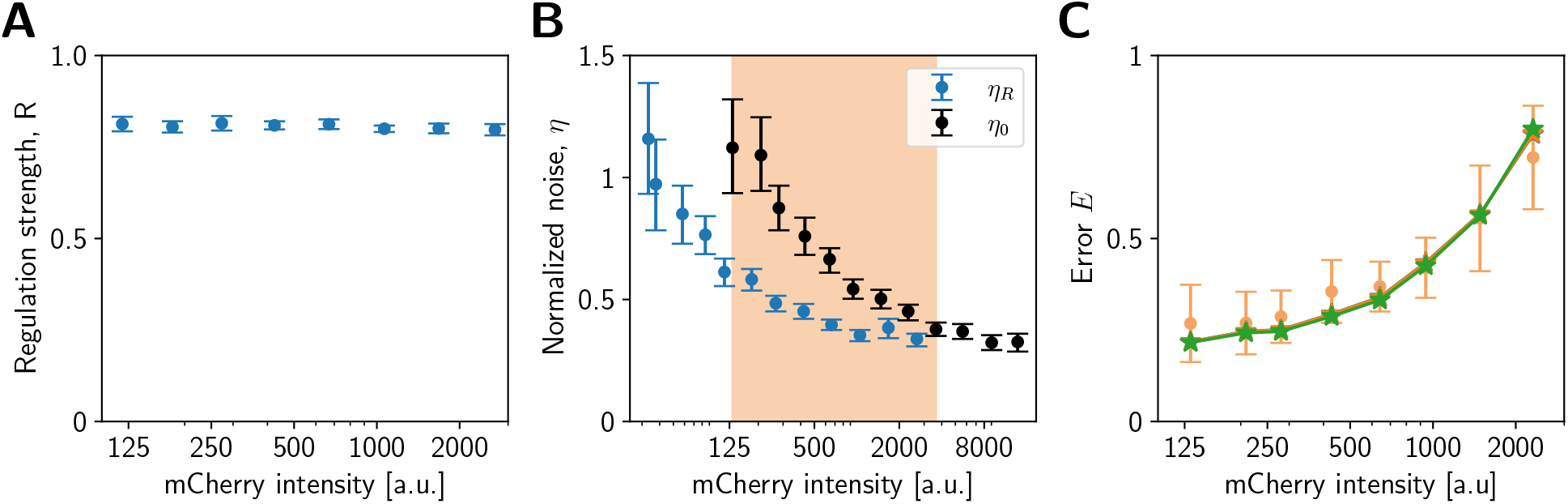
Analysis of the experimental mCherry expression from the *Lats2*-microRNA system [2]. (A) Experimentally determined regulation strengths *R* with varying mCherry intensity. (B) Experimental data for noise levels in unregulated (*η*_0_, black) and regulated (*η*_*R*_, blue) mCherry expression. (C) We span mCherry intensity levels, allowing a direct comparison (within the orange shaded region of (B)) of the noise levels, and transform them to error *E* (orange) using Eq.(S41) and numerical interpolation. The solid lines (multi-color) with stars as points display best-fit curves for *γ*_*μ*_ = 0.001, 0.01, 0.114. These three curves closely overlap, so are indistinguishable.

where the subscript *λ* = 0, *R* applies to unregulated and regulated cases respectively. These are shown in Fig. S3B as blue (regulated data), black (unregulated data). Our theory can be related to the experimental data by estimating the error value,

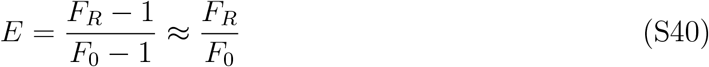

where 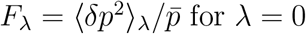, *R* (regulated and unregulated) and the approximation holds when *F*_*R*_, *F*_0_ ≫ 1, which is generally true for protein expression [17]. Thus we can estimate the experimental error *E*^exp^(*X*) from the reported data via

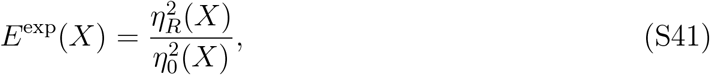

through which we obtain *E*^exp^(*X*) as a function of *X* in the overlapping region (shaded region in Fig. S3B). Since *η*_*R*_(*X*) and *η*_0_(*X*) are not measured at exactly the same *X* values, we use interpolation to match the values, and show the results for the error *E*^exp^(*X*) in Fig. S3C (orange points). To calculate the error bars, we have used the appropriate mean and standard deviation error propagation rules from the original data.

To apply Eq. (S38), we use the average *R*^exp^(*X*) value and perform a chi-squared fitting to *E*^exp^(*X*) using its mean and standard deviations. The quality of fitting is relatively insensitive to *γ*_*μ*_ = *d*_*μ*_*/d*_*mμ*_. Since it is assumed *γ*_*μ*_ should be smaller than 1 (degradation of the miRNA-mRNA complex is faster than miRNA alone), but we do not have good literature estimates for *d*_*mμ*_, we take a biologically plausible range of possible *γ*_*μ*_ values *γ*_*μ*_ = 0.001 − 0.114. The results are shown in Fig. S3C, and curves for different *γ*_*μ*_ values are essentially indistinguishable, all providing good fits to the experimental data.

In Fig. 4A of the main text, we plot the inferred Michaelis-Menten constants normalized by the optimal Michaelis-Menten value, 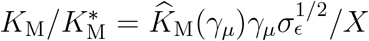, where *σ*_*ϵ*_ is the ratio of metabolic costs of microRNA and mRNA transcripts. The points represent fits using the Table S1 values of *γ*_*μ*_ and *σ*_*ϵ*_, and the colored regions show the uncertainty due to possible ranges of *γ*_*μ*_ and *σ*_*ϵ*_. The ratio 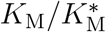 depends very weakly on the choice of *γ*_*μ*_, so the range *γ*_*μ*_ = 0.001 − 0.114 does not significantly affect the estimate (in fact even extending the range to *γ*_*μ*_ *<* 0.001 leaves the uncertainty region for 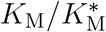 essentially unchanged). For *σ*_*ϵ*_ we chose the range *σ*_*ϵ*_ = 0.0014 − 0.5 as follows. The energetic costs of miRNA and mRNA transcription were introduced in Sec. S.III, with *σ*_*ϵ*_ = *L*_*μ*,pri_*/L*_m,pre_. The average estimated lengths of pri-miRNA *L*_*μ*,pri_ = 2, 500 and pre-mRNAs *L*_m,pre_ = 20, 000 lead to *σ*_*ϵ*_ = 0.125 (Table S1). We can narrow our search knowing that *L*_m,pre_ ≈ 50, 000 for the Lats2 gene [18]. However, the pri-miRNA transcripts are largely unknown due to fast transcription from primary to precursor structure. Moreover, a single pri-miRNA can lead to multiple copies of pre-miRNA. Thus, despite proposed methods [19, 20], we lack this information for mouse microRNAs. Nevertheless, considering the lengths of pre-miRNAs, which are about 70 nucleotides [21], gives an approximate lower bound: *L*_*μ*,pri_ ≥ *L*_*μ*,pre_ = 70, which yields to *σ*_*ϵ*_ = 0.0014 (note that this is a strict underestimation). Since the energetic costs of assembling miRNA are expected to be smaller than mRNA, we use *σ*_*ϵ*_ = 0.5 as an upper bound. The lowest and highest 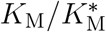 values (boundaries of colored region in Fig. 4A) correspond to the results for lowest and highest values of both *γ*_*μ*_ and *σ*_*ϵ*_.

Finally, as a consistency check, we can attempt to approximately convert the value of the inferred *K*_M_ into units of molar concentration. For this, we need a conversion factor between mCherry intensities and mRNA concentration. In Fig. 4B of Ref. [2], the probability density of mRNA levels in the mESC transcriptome is correlated with the range of mCherry fluorescence intensities. The peak value (most likely mRNA level in RPKM units) corresponds to intensity *X*^*†*^ = 510. While we do not know the peak value for mESC cells in terms of copy numbers, we can use data from mouse fibroblasts [6] as a rough proxy. There the peak occurs around a copy number 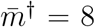(the most likely number of mRNAs from a single gene). Assuming a cell volume of 2,000 *μ*m^3^, this amounts to molar density 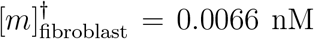. If we take mESC concentration at the peak 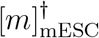 to be similar, then the conversion factor is 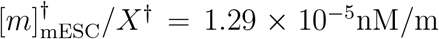 Cherry intensity. We can obtain *K*_M_ in terms of molar concentration using 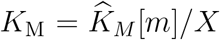 and the conversion factor, which yields *K*_M_ = 8.35, 0.86, 0.1 nM respectively for the possible parameter values *γ*_*μ*_ = 0.001, 0.01, 0.114. For comparison, *K*_M_ for a miRNA system in mouse fibroblasts was measured to be 0.1 nM [15], so our estimated *K*_M_ numbers seem plausible (particularly at *γ*_*μ*_ = 0.114, the value from Table S1 which we used for most of our calculations).

#### miRNA regulation of *sens* in *Drosophila* wing disc cells [16]

These experiments use a similar technique to the one described above, with reporter proteins attached to the *sens* 3’UTR used to investigate the noise regulation mediated by miRNAs. Comparison of noise levels is made between proteins having 3’UTR sequences with and without miR-9a binding sites (mCherry Sens wild type and mCherry Sens mutant). We obtained the regulation strengths *R*^exp^(*X*) by using the raw data for relative expression levels of mCherry Sens wild type and sfGFP Sens mutant proteins. This is plotted in Fig. S4A. The protein expression noise levels are reported in terms of Fano factors *F*_0_ and *F*_*R*_, and we binned the data along different expression levels (Fig. S4B). *E*^exp^ can then be directly obtained from:

**FIG. S4.**
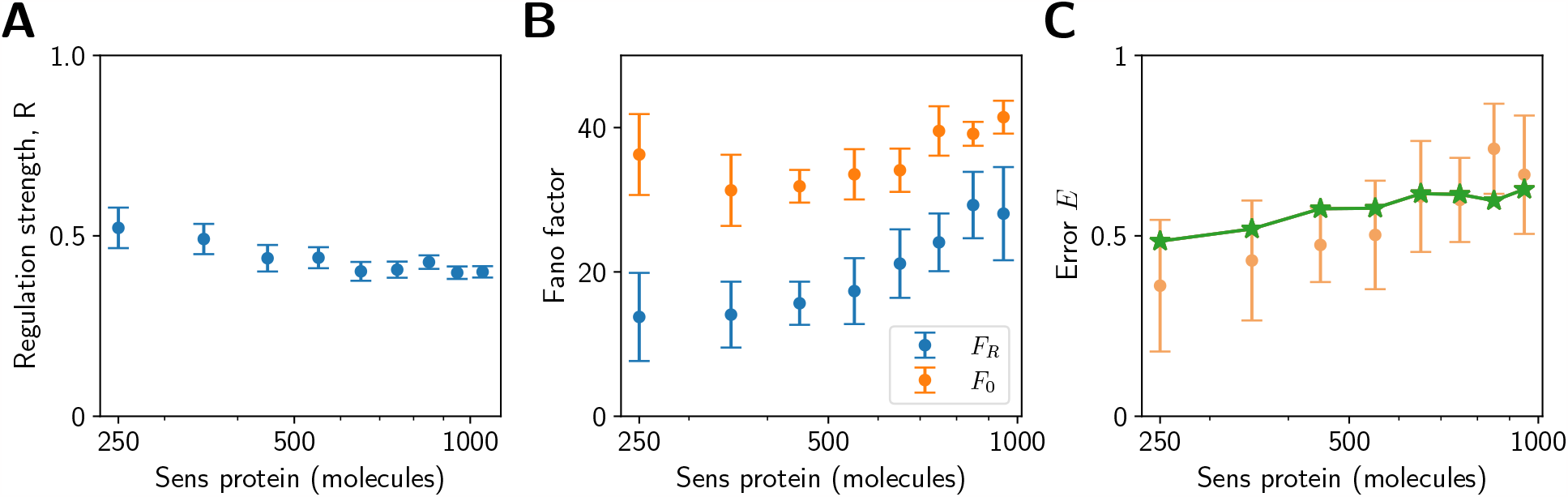
Analysis of the experimental *sens*-miRNA system in *Drosophila* [16]. (A) Experimentally determined regulation strengths *R* with varying Sens protein numbers. (B) Experimental data for the Fano factors of the unregulated (*F*_0_, orange) and regulated (*F*_*R*_, blue) cases. (C) Experimental results for the error *E* (orange), compared to best-fit theoretical curves (multi-color lines with stars correspond to *γ*_*μ*_ = 0.001, 0.01, 0.114, and appear indistinguishable).

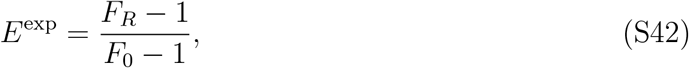

which is shown in Fig. S4C. We then apply a fitting procedure identical to the one described in the *Lats2* example.

Main text Fig. 4B plots the inferred ratio 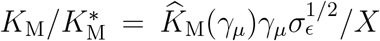, with points representing fits using the parameter values from Table S1. The colored uncertainty region corresponds to varying *γ*_*μ*_ = 0.001 − 0.114 and *σ*_*ϵ*_ = 0.014 − 0.5. In this case, the lower bound for *σ*_*ϵ*_ is obtained using the pre-mRNA length of the *sens* gene *L*_m,pre_ ≈ 5, 000 [22] and *L*_*μ*,pri_ ≥ *L*_*μ*,pre_ = 70, yielding *σ*_*ϵ*_ = 0.014.

## References

[1] X. Yang, M. Heinemann, J. Howard, G. Huber, S. Iyer-Biswas, G. Le Treut, M. Lynch, K. L. Montooth, D. J. Needleman, S. Pigolotti, et al., Physical bioenergetics: Energy fluxes, budgets, and constraints in cells, Proc. Natl. Acad. Sci. 118, e2026786118 (2021).

[2] K. Chen and N. Rajewsky, The evolution of gene regulation by transcription factors and micrornas, Nat. Rev. Genet. 8, 93 (2007).

[3] D. P. Bartel, Metazoan micrornas, Cell 173, 20 (2018).

[4] M. Osella, C. Bosia, D. Corá, and M. Caselle, The role of incoherent microrna-mediated feedforward loops in noise buffering, PLoS Comput. Biol. 7, e1001101 (2011).

[5] J. M. Schmiedel, S. L. Klemm, Y. Zheng, A. Sahay, N. Blüthgen, D. S. Marks, and A. van Oudenaarden, Microrna control of protein expression noise, Science 348, 128 (2015).

[6] V. Siciliano, I. Garzilli, C. Fracassi, S. Criscuolo, S. Ventre, and D. Di Bernardo, Mirnas confer phenotypic robustness to gene networks by suppressing biological noise, Nat. Comm. 4, 2364 (2013).

[7] E. Hornstein and N. Shomron, Canalization of develop-ment by micrornas, Nat. Genet. 38, S20 (2006).

[8] M. S. Ebert and P. A. Sharp, Roles for micrornas in conferring robustness to biological processes, Cell 149, 515 (2012).

[9] K. J. Peterson, M. R. Dietrich, and M. A. McPeek, Micrornas and metazoan macroevolution: insights into canalization, complexity, and the cambrian explosion, Bioessays 31, 736 (2009).

[10] C. Alberti and L. Cochella, A framework for understanding the roles of mirnas in animal development, Development 144, 2548 (2017).

[11] D. T. Gillespie, The chemical langevin equation, J. Chem. Phys. 113, 297 (2000).

[12] G. Tkačik and A. M. Walczak, Information transmis-sion in genetic regulatory networks: a review, Journal of Physics: Condensed Matter 23, 153102 (2011).

[13] J. Tsang, J. Zhu, and A. Van Oudenaarden, Micrornamediated feedback and feedforward loops are recurrent network motifs in mammals, Mol. Cell 26, 753 (2007).

[14] N. J. Martinez and A. J. Walhout, The interplay between transcription factors and micrornas in genome-scale regulatory networks, Bioessays 31, 435 (2009).

[15] D. Baek, J. Villén, C. Shin, F. D. Camargo, S. P. Gygi, and D. P. Bartel, The impact of micrornas on protein output, Nature 455, 64 (2008).

[16] M. Selbach, B. Schwanhäusser, N. Thierfelder, Z. Fang, R. Khanin, and N. Rajewsky, Widespread changes in protein synthesis induced by micrornas, Nature 455, 58 (2008).

[17] D. Hathcock, J. Sheehy, C. Weisenberger, E. Ilker, and M. Hinczewski, Noise filtering and prediction in biological signaling networks, IEEE Trans. Mol. Biol. Multi-Scale Commun. 2, 16 (2016).

[18] M. Lynch and G. K. Marinov, The bioenergetic costs of a gene, Proc. Natl. Acad. Sci. 112, 15690 (2015).

[19] L. M. Wee, C. F. Flores-Jasso, W. E. Salomon, and P. D. Zamore, Argonaute divides its rna guide into domains with distinct functions and rna-binding properties, Cell 151, 1055 (2012).

[20] M. Del Giudice, S. Bo, S. Grigolon, and C. Bosia, On the role of extrinsic noise in microrna-mediated bimodal gene expression, PLoS Comput. Biol. 14, e1006063 (2018).

[21] E. Ferro, C. E. Bena, S. Grigolon, and C. Bosia, microrna-mediated noise processing in cells: A fight or a game?, Comput. Struct. Biotechnol. J. 18, 642 (2020).

[22] R. Lorenz, S. H. Bernhart, C. Höner zu Siederdissen, H. Tafer, C. Flamm, P. F. Stadler, and I. L. Hofacker, Viennarna package 2.0, Algorithms Mol. Biol. 6, 1 (2011).

[23] D. C. Ellwanger, F. A. Büttner, H.-W. Mewes, and V. Stümpflen, The sufficient minimal set of mirna seed types, Bioinformatics 27, 1346 (2011).

[24] N. Wiener, Extrapolation, interpolation, and smoothing of stationary time series, Vol. 2 (MIT press Cambridge, 1949).

[25] A. N. Kolmogorov, Interpolation and extrapolation of stationary random sequences, Izv. Akad. Nauk SSSR Ser. Mat. 5, 3 (1941).

[26] H. W. Bode and C. E. Shannon, A simplified derivation of linear least square smoothing and prediction theory, Proc. Inst. Radio. Engin. 38, 417 (1950).

[27] M. Hinczewski and D. Thirumalai, Cellular signaling networks function as generalized wiener-kolmogorov filters to suppress noise, Phys. Rev. X 4, 041017 (2014).

[28] N. B. Becker, A. Mugler, and P. R. ten Wolde, Optimal prediction by cellular signaling networks, Phys. Rev. Lett. 115, 258103 (2015).

[29] M. Hinczewski and D. Thirumalai, Noise control in gene regulatory networks with negative feedback, J. Phys. Chem. B (2016).

[30] D. Hartich and U. Seifert, Optimal inference strategies and their implications for the linear noise approximation, Phys. Rev. E 94, 042416 (2016).

[31] P. R. ten Wolde, N. B. Becker, T. E. Ouldridge, and A. Mugler, Fundamental limits to cellular sensing, J. Stat. Phys. 162, 1395 (2016).

[32] C. Zechner, G. Seelig, M. Rullan, and M. Khammash, Molecular circuits for dynamic noise filtering, Proc. Natl. Acad. Sci. U. S. A. 113, 4729 (2016).

[33] T.-L. Wang, B. Kuznets-Speck, J. Broderick, and M. Hinczewski, The price of a bit: energetic costs and the evolution of cellular signaling, bioRxiv, 2020.10.06.327700 (2020).

[34] W. Sung, M. S. Ackerman, S. F. Miller, T. G. Doak, and M. Lynch, Drift-barrier hypothesis and mutationrate evolution, Proc. Natl. Acad. Sci. 109, 18488 (2012).

[35] J. H. Gillespie, Population Genetics: A Concise Guide (JHU Press, 2010).

[36] B. Charlesworth, Effective population size and patterns of molecular evolution and variation, Nat. Rev. Genet. 10, 195 (2009).

[37] M. Kimura, The neutral theory of molecular evolution (Cambridge University Press, 1983).

[38] E. Ilker and M. Hinczewski, Modeling the growth of organisms validates a general relation between metabolic costs and natural selection, Phys. Rev. Lett. 122, 238101 (2019).

[39] R. Giri, D. K. Papadopoulos, D. M. Posadas, H. K. Potluri, P. Tomancak, M. Mani, and R. W. Carthew, Ordered patterning of the sensory system is susceptible to stochastic features of gene expression, eLife 9, e53638 (2020).

[40] N. Yabuta, N. Okada, A. Ito, T. Hosomi, S. Nishihara, Y. Sasayama, A. Fujimori, D. Okuzaki, H. Zhao, M. Ikawa, et al., Lats2 is an essential mitotic regulator required for the coordination of cell division, Journal of Biological Chemistry 282, 19259 (2007).

[41] N. Furth and Y. Aylon, The lats1 and lats2 tumor suppressors: beyond the hippo pathway, Cell Death & Differentiation 24, 1488 (2017).

[42] F. J. Navarro and D. C. Baulcombe, mirna-mediated regulation of synthetic gene circuits in the green alga chlamydomonas reinhardtii, ACS Synth. Biol. 8, 358 (2019).

[43] A. Kotowska-Zimmer, M. Pewinska, and M. Olejniczak, Artificial mirnas as therapeutic tools: Challenges and opportunities, Wiley Interdiscip. Rev. RNA 12, e1640 (2021).

[44] T. Kang, T. Quarton, C. M. Nowak, K. Ehrhardt, A. Singh, Y. Li, and L. Bleris, Robust filtering and noise suppression in intragenic mirna-mediated host regula-tion, iScience 23 (2020).

[45] L. Wei, S. Li, P. Zhang, T. Hu, M. Q. Zhang, Z. Xie, and X. Wang, Characterizing microrna-mediated modulation of gene expression noise and its effect on synthetic gene circuits, Cell Rep. 36 (2021).

[46] X.-J. Tian, H. Zhang, J. Zhang, and J. Xing, Reciprocal regulation between mrna and microrna enables a bistable switch that directs cell fate decisions, FEBS Lett. 590, 3443 (2016).

[47] X. Li, J. J. Cassidy, C. A. Reinke, S. Fischboeck, and R. W. Carthew, A microrna imparts robustness against environmental fluctuation during development, Cell 137, 273 (2009).

[48] E. M. Ozbudak, M. Thattai, I. Kurtser, A. D. Grossman, and A. Van Oudenaarden, Regulation of noise in the expression of a single gene, Nat. Genet. 31, 69 (2002).

[49] A. Raj and A. van Oudenaarden, Nature, nurture, or chance: stochastic gene expression and its consequences, Cell 135, 216 (2008).

[50] R. Fan and A. Hilfinger, The effect of microrna on protein variability and gene expression fidelity, Biophys. J. 122 (2023).

[51] O. S. Rissland, A. O. Subtelny, M. Wang, A. Lugowski, B. Nicholson, J. D. Laver, S. S. Sidhu, C. A. Smibert, H. D. Lipshitz, and D. P. Bartel, The influence of micror-nas and poly (a) tail length on endogenous mrna–protein complexes, Genome Biol. 18, 1 (2017).

[52] M. Jens and N. Rajewsky, Competition between target sites of regulators shapes post-transcriptional gene regulation, Nat. Rev. Genet. 16, 113 (2015).

[53] R. L. Skalsky and B. R. Cullen, Viruses, micrornas, and host interactions, Annu. Rev. Microbiol. 64, 123 (2010).

[54] G. Mahmoudabadi, R. Milo, and R. Phillips, Energetic cost of building a virus, Proc. Natl. Acad. Sci. 114, E4324 (2017).

## References

[1] D. T. Gillespie, J. Chem. Phys. 113, 297 (2000).

[2] J. M. Schmiedel, S. L. Klemm, Y. Zheng, A. Sahay, N. Blüthgen, D. S. Marks, and A. van Oudenaarden, Science 348, 128 (2015).

[3] S. Mukherji, M. S. Ebert, G. X. Zheng, J. S. Tsang, P. A. Sharp, and A. Van Oudenaarden, Nat. Genet. 43, 854 (2011).

[4] M. Lynch and G. K. Marinov, Proc. Natl. Acad. Sci. 112, 15690 (2015).

[5] D. Hathcock, J. Sheehy, C. Weisenberger, E. Ilker, and M. Hinczewski, IEEE Trans. Mol. Biol. Multi-Scale Commun. 2, 16 (2016).

[6] B. Schwanhäusser, D. Busse, N. Li, G. Dittmar, J. Schuchhardt, J. Wolf, W. Chen, and M. Selbach, Nature 473, 337 (2011).

[7] M. J. Marzi, F. Ghini, B. Cerruti, S. De Pretis, P. Bonetti, C. Giacomelli, M. M. Gorski, T. Kress, M. Pelizzola, H. Muller, et al., Genome research 26, 554 (2016).

[8] E. R. Kingston and D. P. Bartel, Genome Res. 29, 1777 (2019).

[9] B. Alberts, A. Johnson, J. Lewis, D. Morgan, M. Raff, K. Roberts, and P. Walter, Molecular Biology of the Cell (Garland Science, 2015).

[10] G. Song and L. Wang, PloS One 3, e3574 (2008).

[11] H. K. Saini, A. J. Enright, and S. Griffiths-Jones, BMC Genom. 9, 1 (2008).

[12] R. Milo, P. Jorgensen, U. Moran, G. Weber, and M. Springer, Nucleic Acids Res. 38, D750 (2010), http://bionumbers.hms.harvard.edu/.

[13] R. Lorenz, S. H. Bernhart, C. Höner zu Siederdissen, H. Tafer, C. Flamm, P. F. Stadler, and L. Hofacker, Algorithms Mol. Biol. 6, 1 (2011).

[14] V. Agarwal, G. W. Bell, J.-W. Nam, and D. P. Bartel, eLife 4, e05005 (2015).

[15] L. M. Wee, C. F. Flores-Jasso, W. E. Salomon, and P. D. Zamore, Cell 151, 1055 (2012).

[16] R. Giri, D. K. Papadopoulos, D. M. Posadas, H. K. Potluri, P. Tomancak, M. Mani, and R. W. Carthew, eLife 9, e53638 (2020).

[17] E. M. Ozbudak, M. Thattai, I. Kurtser, A. D. Grossman, and A. Van Oudenaarden, Nature genetics 31, 69 (2002).

[18] J. A. Blake, R. Baldarelli, J. A. Kadin, J. E. Richardson, C. L. Smith, and C. J. Bult, Nucl. Acids Res. 49, D981 (2021).

[19] J. Qian, Z. Zhang, J. Liang, Q. Ge, X. Duan, F. Ma, and F. Li, Genomics 97, 294 (2011).

[20] K. Bedi, M. T. Paulsen, T. E. Wilson, and M. Ljungman, NAR genomics and bioinformatics 2, lqz014 (2020).

[21] Y. Lee, K. Jeon, J.-T. Lee, S. Kim, and V. N. Kim, The EMBO journal 21, 4663 (2002).

[22] L. S. Gramates, J. Agapite, H. Attrill, B. R. Calvi, M. A. Crosby, G. Dos Santos, J. L. Goodman, D. Goutte-Gattat, V. K. Jenkins, T. Kaufman, et al., Genetics 220, iyac035 (2022).

